# Stable Hrd1 tetramers at the heart of the retrotranslocon in living cells

**DOI:** 10.64898/2026.06.16.732623

**Authors:** Tim Abel, Shubha Subramanya, Brandon M. Liu, Anja Schütz, Jennifer Lippincott-Schwartz, Sonya Neal, Thomas Sommer, Christopher Obara

## Abstract

Terminally misfolded or damaged proteins in the endoplasmic reticulum (ER) are degraded by the proteasome in a process termed endoplasmic reticulum-associated degradation (ERAD). To reach the proteasome in the cytosol, cargo proteins must be dislocated across the ER lipid bilayer. Retrograde transport is performed by a set of membrane-spanning multi-protein complexes that are notoriously difficult to study. Despite decades of genetic and biochemical analysis, there is little consensus regarding its molecular mechanism. Here, we characterize the dynamic assembly of the mammalian Hrd1 complex, one of the most thoroughly studied mammalian dislocons, using in situ multicolor single-particle tracking. Surprisingly, quantitative dual-color tracking reveals that the majority of Hrd1 is assembled into stable homo-tetramers in situ. Using a herein developed single-molecule assay based on binding competition, we show that this tetramerization is driven by a short stretch (Hrd1_479-530_, HAF-H) within the cytosolic domain of Hrd1. Combining classical purification assays, quantitative imaging, and structural predictions, we demonstrate that Hrd1_479-530_ forms a highly stable tetrameric helix bundle via a conserved hydrophobic coiled-coil motif. While higher order assemblies of Hrd1 have been previously implicated, this work offers direct evidence of their role in dislocation, demonstrating that quantitative single-molecule imaging can yield information on species previously hidden from classical biochemistry.

## Introduction

The endoplasmic reticulum (ER) is a single, continuous, membrane-enclosed compartment and the birthplace of nearly all membrane and secreted proteins.^1,2^ The error-prone nature of protein folding challenges the ER with a constant load of proteins that never obtain a functional fold, encompassing diverse protein topologies and sizes. To prevent accumulation of these potentially harmful species, misfolded proteins need to be efficiently detected and disposed of. In the ER, targeted protein degradation is realized by a process termed endoplasmic reticulum-associated degradation (ERAD). ERAD detects misfolded proteins and transports them to the cytosol where they are degraded by the proteasome.^3^

Membrane-associated ubiquitin-E3-ligase complexes form the core of the ERAD system. These protein complexes mediate both a substrate’s ER-to-cytosol transport, referred to as retro-translocation, as well as its subsequent conjugation to ubiquitin, a post-translational modification flagging them for degradation. The conjugation of polyubiquitin chains acts as the signal for the recruitment of the cytosolic AAA ATPase VCP/p97, which catalyzes the extraction of the misfolded protein from the ER and routes it to the 26S proteasome for degradation.^4^ Of the at least 10 different ERAD E3-ligase complexes identified in humans,^5^ the complex surrounding the ubiquitin-ligase Hrd1 is the best characterized. Hrd1 targets a surprisingly broad spectrum of proteins, including both luminal-soluble and ER membrane-integrated proteins.^6,7^ While its role in the ubiquitination of substrates is well understood, the mechanisms underlying the retro-translocation of client proteins remain enigmatic. The heterogeneous cargo that Hrd1 needs to transport across the hydrophobic ER bilayer includes bulky glycan modifications^8,9^ and potentially even partially folded domains.^10–12^ Currently, the biochemical and structural data available do not support a conclusive mechanism for retro-translocation. Thus, a clear picture of the architecture and dynamics of the complex will empower speculation on the mechanism of dislocation.

Hrd1 assembles in a large multicomponent transmembrane protein complex, a protein class that remains a challenge even for modern biochemical analysis. The Hrd1 complex has been extensively studied in both yeast and mammals, both structurally and functionally. Presumably, all its components have been identified^13^ and most were assigned a function. While several inconsistent 3D structures have been generated,^14–18^ a functional holo-complex has not been successfully purified and characterized. Purification attempts have largely been challenged by high variance in both composition and overall size of isolated complexes.^17,19–21^ While the critical components have been identified, there is currently no consensus on the structural arrangement or even the stoichiometry of the Hrd1 complex. Whether the dynamic behavior in biochemical experiments is a biological feature of the complex or a result of the experimental challenges remains to be explored.

Live-cell fluorescence imaging offers an opportunity to directly observe molecular machinery at its site of function. In this study, we employ fluorescent dual-color single-molecule tracking (dcSMT) to interrogate the stoichiometry and architecture of the mammalian Hrd1 complex in its native environment. While recent work has demonstrated the value of dcSMT in tracking protein complex dynamics,^22,23^ its potential for biochemical characterization of protein complexes remains largely untapped. Here, we introduce a series of correlative dcSMT assays, which we use to show that Hrd1 forms a highly stable homo-tetramer via its cytosolic HAF-H domain. Combining these results with structural predictions and classical domain truncation experiments, we show that higher-order oligomeric Hrd1 is the main assembly form of Hrd1.

## Results

### Single-Molecule Localization and Dynamics of Endogenous Hrd1-Halo

Single-molecule imaging requires labeling the protein of interest with bright and photostable fluorophore. HaloTag is an engineered self-labeling protein tag that covalently and specifically binds its ligand, which in turn can be conjugated to a fluorophore of choice.^24^ Fused to the protein of interest, HaloTag enables labeling with bright chemical dyes compatible with single-molecule imaging. Our attempts to tag Hrd1 C-terminally resulted in a non-functional form (data not shown), presumably due to the tag masking the C-terminal VCP interaction motif of Hrd1. Bioinformatic analysis of Hrd1 identified a highly disordered and evolutionary little conserved stretch without apparent functional relevance located between the cytosolic HAF-H and the C-terminal VCP interaction domain.^21^ Using this to our advantage, we integrated the HaloTag at this position (downstream of Gly588) flanked by short flexible peptide linkers (Figure 1A).

**Figure 1:**
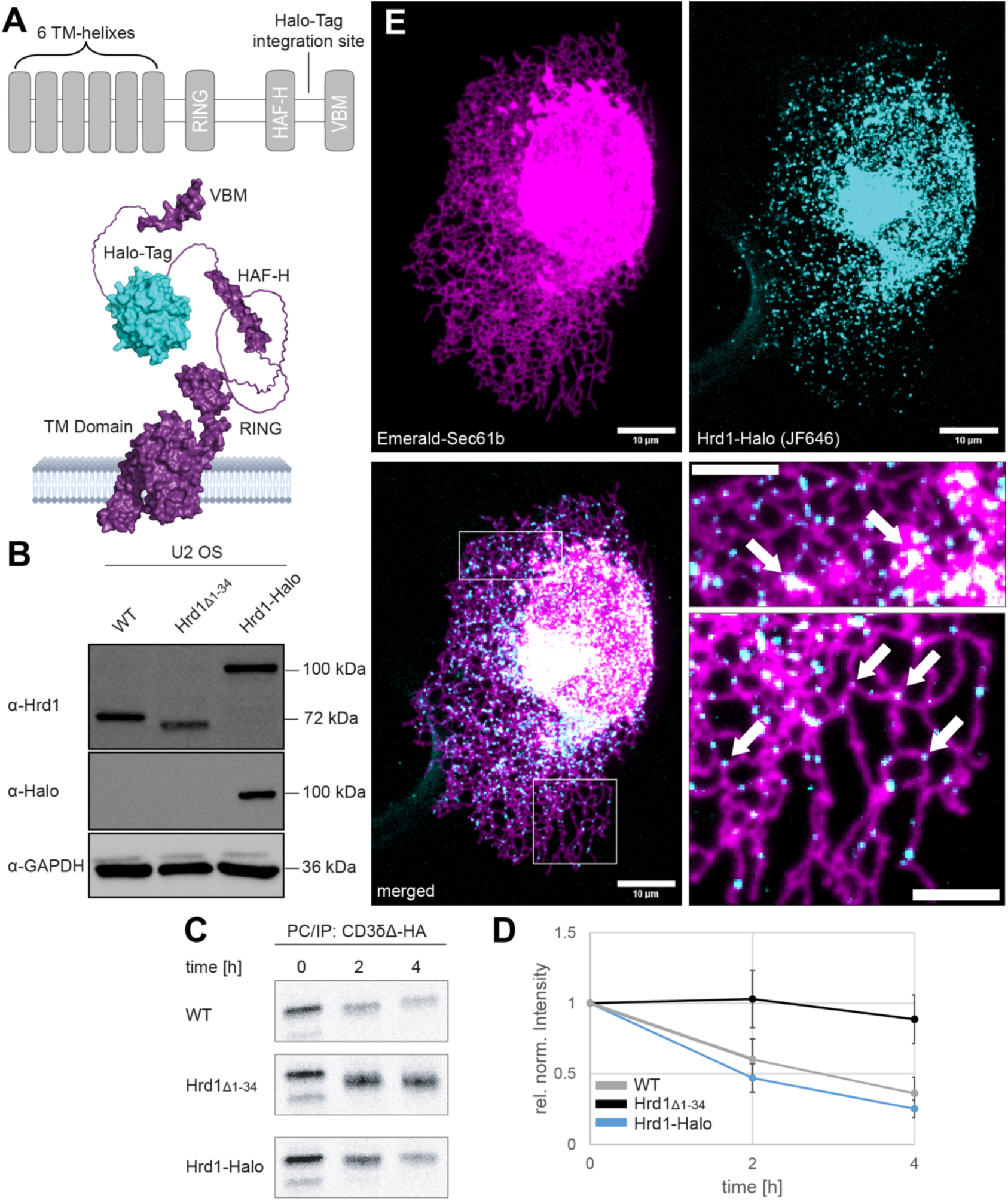
Transgenic U-2 OS Hrd1-Halo cells display unaltered behavior in Hrd1-mediated degradation and complex formation. **A)** Hrd1 domain architecture schematic visualizing the internal HaloTag placement **B)** Western blot of cell lysates from genetically modified U-2 OS cell lines. **C)** Autoradiogram of the Hrd1-substrate CD3-δΔ-HA after pulse-chase (PC) labeling with 35S Cys/Meth and immunoprecipitation (IP) using α-HA IgG. Samples were taken at indicated timepoints and separated using SDS-PAGE. **D)** Quantification of C (biological triplicate). Band intensities were normalized to t = 0. **E)** Spinning-Disk images of fixed U-2 OS Hrd1-Halo. Cells were transfected with Emerald-Sec61b as an ER-marker and stained with JF646-Halo. Scalebar, 4 µm in magnifications.

To prevent artifacts resulting from overexpression,^20^ we integrated the HaloTag sequence into the genomic locus of Hrd1. We employed targeted CRISPR/Cas9 gene editing to generate U-2 OS cells expressing chimeric Hrd1-Halo directly from its endogenous genomic locus (Figure 1B). The resulting monoclonal cell line Hrd1-Halo was then characterized for its activity in ERAD. To this end, we followed the degradation of the model Hrd1-targeted substrate CD3-δΔ-HA^6^ using radioactive pulse-chase assays and analyzed the composition of the Hrd1-complex via western blotting and pull-down experiments (Figure 1C/D, S1).

CD3-δΔ-HA was rapidly degraded in U-2 OS cells but not in cells expressing a dysfunctional truncated version of Hrd1 (Hrd1Δ1-34). In the endogenously tagged Hrd1-Halo cells, CD3-δΔ-HA was degraded with a half-life comparable to the one in wild type cells, demonstrating that the insertion of the HaloTag did not detectibly interfere with the function or kinetics of Hrd1 in mediating protein degradation. Consistent with this and previous literature, the abundance of Hrd1-dependent interaction partners Sel1L and HERPUD1 was unchanged in comparison to wild type cells, whereas they were substantially increased upon loss of Hrd1 function in Hrd1Δ1-34 cells (Figure S1).^25,26^ Taken together, the complex formed around Hrd1-Halo retains its activity and the expression level of the associated components remains unchanged.

Genomic labeling of Hrd1 with the HaloTag offers a direct view into the formation of the endogenous Hrd1 complex using fluorescence microscopy. Initially, we assessed the subcellular distribution of Hrd1-Halo with confocal microscopy. Briefly, Hrd1-Halo cells were transiently transfected with a construct for the expression mEmerald-Sec61 as an ER marker, labeled with a fluorescent JF646-conjugated HaloTag ligand,^27^ and imaged using confocal spinning disk microscopy (Figure 1E). As expected, the highest density of Hrd1-Halo was observed in the perinuclear ER around the nuclear envelope. In less dense ER regions, Hrd1-Halo appeared as spatially-isolated, diffraction-limited spots. While Hrd1-Halo was localized to all ER topologies, it appeared to be enriched in ER three-way junctions and sheets (Figure 1E, magnified). Interestingly, this preference to membrane domains traditionally associated with low membrane curvature has previously been observed for other macromolecular protein complexes in the ER.^28,29^

When the peripheral ER of JF646-labeled Hrd1-Halo cells was imaged using high-speed, single-color SMT, Hrd1-Halo was detectable as individually resolvable diffraction-limited punctae (Figure 2A, <u>Video S1</u>). These spots exhibited discrete bleaching steps, indicating that most puncta contained a small number of individual Hrd1-molecules diffusing through the ER together (Figure 2B, <u>Video S2</u>). To analyze the dynamic properties of these complexes, we localized and tracked single spots over time using the Fiji plugin Trackmate.^30^ Motion of protein complexes in the ER membrane has been suggested to be largely diffusion-dominated, the rate of which is directly related to the size-dependent drag imposed by the membrane.^31,32^ To quantify Hrd1 complex motion, we employed a modified form of a classical mean squared displacement (MSD)-fit. Fitting the 2D-MSD of 1582 single-particle trajectories yielded an average diffusion coefficient of 0.033 ± 0.057 µm^2^/s, notably smaller than the coefficient described for many other proteins localized to the ER membrane (Figure 2C).^33–35^

**Figure 2:**
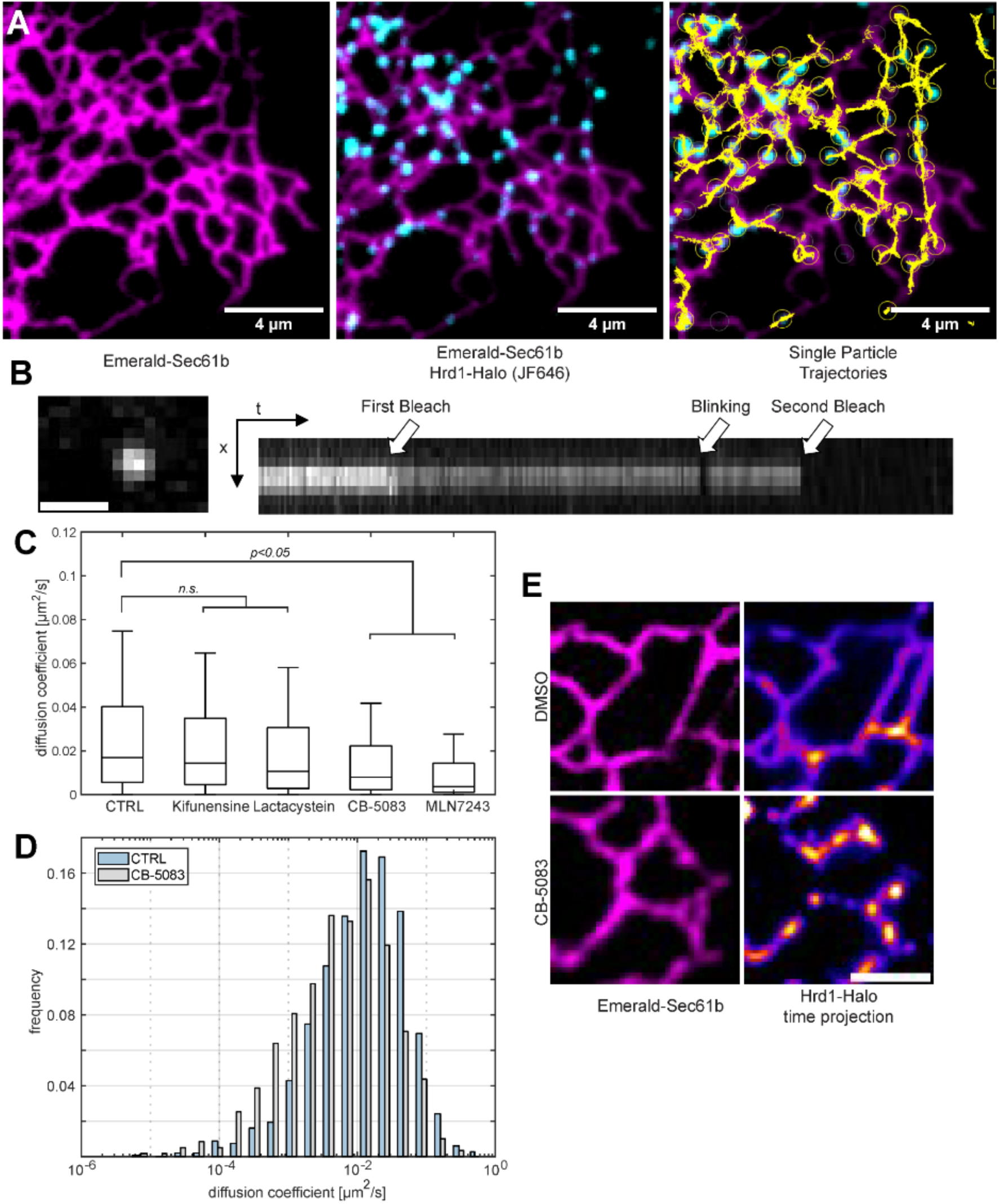
Hrd1-Halo can be visualized and tracked with single complex resolution and its mobility is altered during substrate processing. **A)** Total-Internal-Reflection (TIRF) SMT of U-2 OS Hrd1-Halo cells stained with JF646-Halo and transfected with Emerald-Sec61b. Single Particle Trajectories were generated by tracking diffraction-limited Hrd1-Halo punctae and plotted onto the original data. **B)** Representative single Hrd1-Halo complex as observed in SMT (scalebar, 2 µm). The kymograph of the stabilized recording shows discrete bleaching events. **C)** Mean diffusion coefficients extracted from Hrd1-Halo trajectories after prolonged chemical inhibitions (6h) targeting various steps in ERAD. n(cells) = 66/38/23/36/20 n(trajectories) = 1582/958/495/643/416. **D)** Comparison of the histograms of Hrd1-Halo diffusion coefficients between untreated and p97-inhibited (CB-5083). n(cells) = 66/36 n(trajectories) = 1582/643. **E)** Comparison of representative TIRF images of U-2 OS Hrd1-Halo cells after inhibition of p97 (CB-5083). For visualization the single molecule recordings of Hrd1-Halo were summed across time. Scalebar, 4 µm.

Biochemical approaches have demonstrated changes in the size and composition of the Hrd1 complex in response to chemical inhibition of the ubiquitin-proteasome system.^20^ We predicted that these changes might be reflected in the diffusive properties of the complex. We therefore treated Hrd1-Halo cells with different specific ERAD-inhibitors and monitored their effects on Hrd1-Halo diffusion (Figure 2C). When inhibiting steps up- or down-stream of dislocation such as by inhibiting the proteasome (Lactacystein) or deglycosylation (Kifunensine), there was no effect on the average effective diffusion coefficient. However, inhibiting p97 (CB-5083) or ubiquitination (MLN7243) and thereby substrate extraction from the ER caused a significant reduction of the average effective diffusion coefficient (0.02 ± 0.034 µm^2^/s and 0.015 ± 0.035 µm^2^/s, respectively). Additionally, we observed a relative enrichment of highly restricted punctae (Figure 2D) that appear to have a preference for low curvature domains (Figure 2E, <u>Video S3</u>). Interestingly, both inhibitors are thought to affect ERAD by stalling the retro-translocation of substrates. Thus, the slow-moving Hrd1-Halo particles might represent fully assembled Hrd1 holo-complexes engaged in transporting client proteins out of the ER, but more conclusive data will be needed to validate this in future studies.

### Quantitative Dual-Color Tracking of Hrd1-Halo

Both the low diffusion coefficient and the multiple discrete bleaching steps of Hrd1-Halo punctae (Figure 2B/C) suggest an organization of Hrd1-Halo as higher-order multimeric complexes. We next sought to investigate this issue in more detail and extended the SPT imaging approach using a dual-color setup. We took advantage of the availability of distinctly colored fluorochromes conjugated to the HaloTag ligand. Thus, HaloTag multimers, when simultaneously labeled using two different ligands, are observable in the form of colocalized diffraction-limited spots in each color exhibiting correlated movement (Figure 3A).

**Figure 3:**
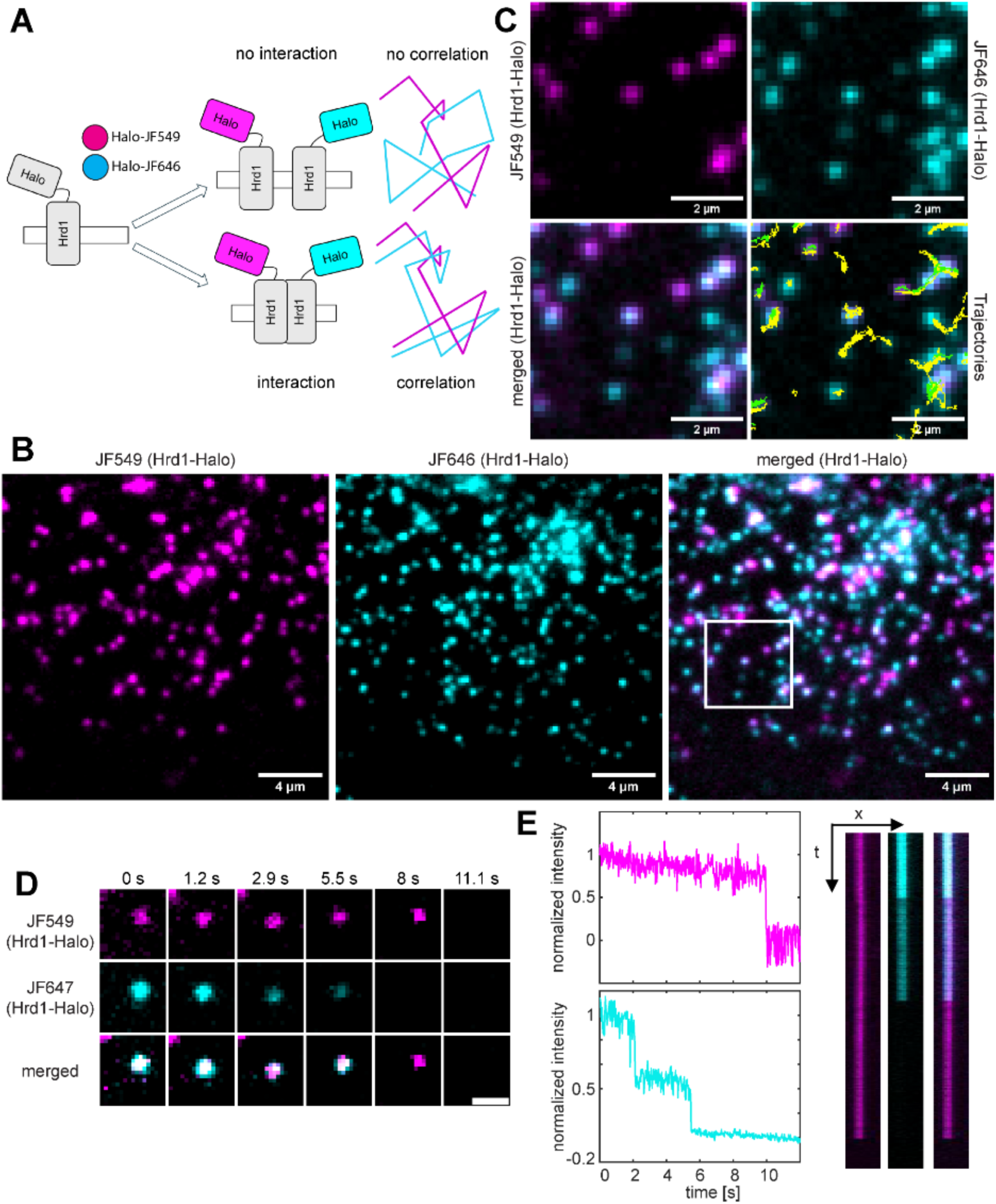
Hrd1-Halo oligomers can readily be observed by TIRF and mixed-dye labeling. **A)** Schematic of mixed-dye labeling of Hrd1-Halo. **B)** Dual-color SMT of U-2 OS Hrd1-Halo cells labeled with JF549-Halo (250 nM) and JF647-Halo (1 µM). **C)** Close-up of the area magnified in (B) together with the generated single molecule trajectories (green = JF549, yellow = JF646). **D)** Representative recordings of a Hrd1-Halo complex demonstrating correlated movement in dcSMT and discrete bleaching. Scalebar, 1 µm. **E)** Single molecule intensity traces and kymographs of (D). The intensity values were baseline corrected and normalized to t=0.

The majority of these correlated trajectories showed photobleaching indicative of two emitters: a single emitter bleaching in one step in each channel. However, due to incomplete labeling of the HaloTag, this is consistent with a higher order oligomer, and not conclusive for a dimer. Accordingly, in an estimated 5% of all cases, we observed an additional discrete bleaching step in one of the two colors in a correlated trajectory pair (Figure 3D/E, <u>Video S5</u>), showing that a significant subset of complexes contains more than two fluorophores and thus must have more than two copies of Hrd1-Halo.

In theory, the percentage of correlated trajectories should allow us to draw direct conclusions on the degree of oligomerization of the protein of interest. Conjugation of dyes to the HaloTag is expected to follow a stochastic distribution (Figure 4A) and, in consequence, higher-order oligomers should more likely contain differently labeled subunits. In practice, experimental hurdles such as incomplete labeling or undetected events interfere with a direct derivation of the oligomeric state just from the occurrence of differently labeled sub-populations alone. The probability of observing correlation for a given oligomer only depends on 1) the ratio of labeling between the both dyes and 2) the efficiency of labeling and detecting the Halo-tag.

**Figure 4:**
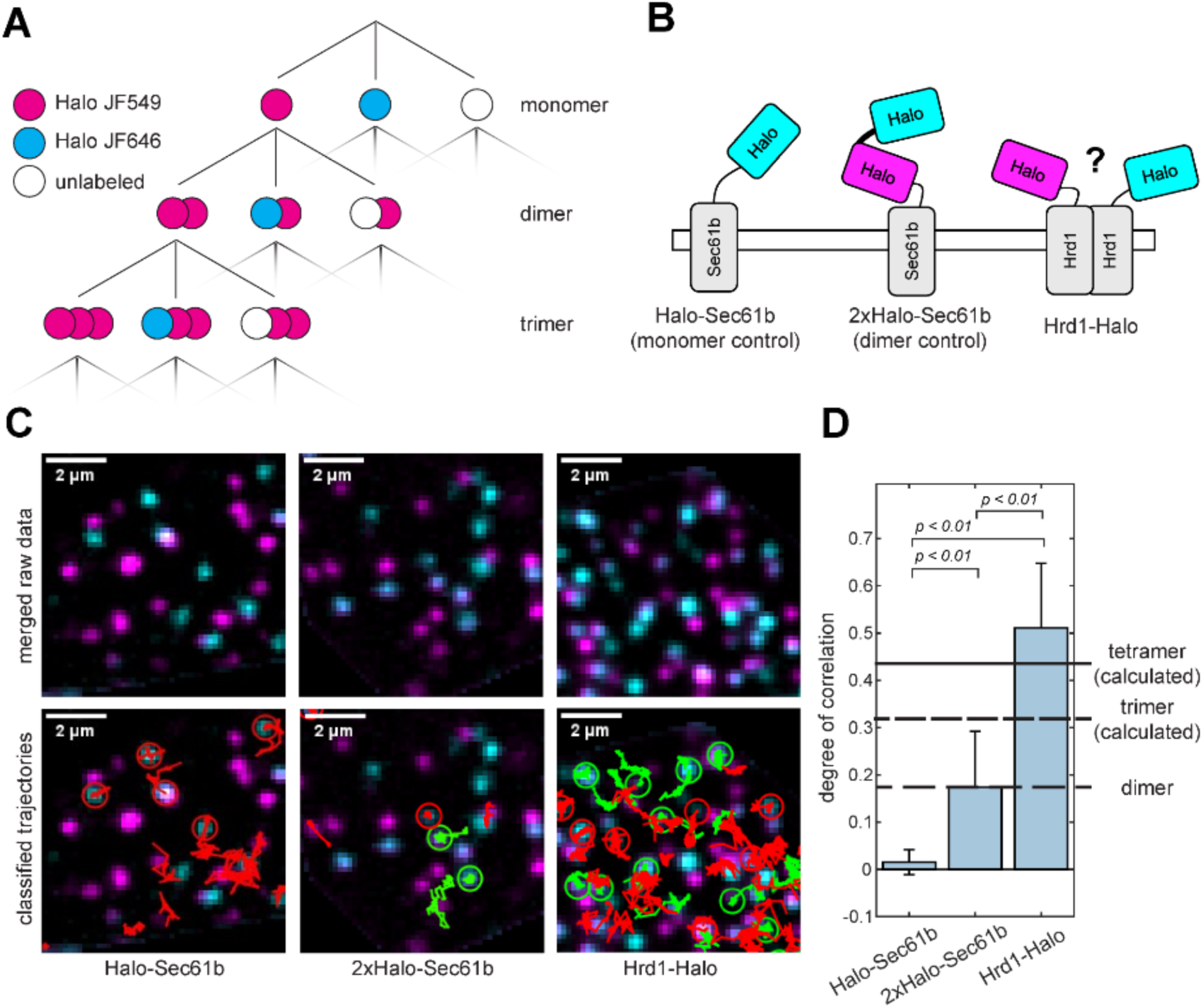
Quantification of correlated movement shows that Hrd1-Halo assembles in multimers containing more than two copies. **A)** Multinomial distribution of potential labeling states as a result of stochastic mixed-dye labeling. **B)** Schematic representations of Halo-Sec61b and 2x-Halo-Sec61b used for comparison in quantitative dcSMT. **C)** Representative SMT recordings and trajectories of the constructs depicted in B after mixed-dye labeling. The trajectories were color-coded regarding the correlation between both channels (green = trajectory with correlation partner, red = uncorrelated). Constructs coding for Halo-Sec61b and 2xHalo-Sec61b were lentivirally transduced and imaged after 6 h to reach appropriate densities. **D)** Cellwise Mean degree of correlation between the constructs shown in B. Correlation was detected by prolonged proximity (7 frames, 150 nm) of single emitters localized with sub-pixel accuracy. The values for tri- and tetramers were calculated, assuming the multinomial distribution shown in (A) using labeling parameters extracted from the Halo-Sec61b and 2xHalo-Sec61b datasets. A one-sided ANOVA was used to test statistical significance. n(cells) = 31/33/66, n(trajectories) = 884/1192/3082.

To address this issue, we aimed to compare the fraction of correlated trajectories of mixed-labeled Hrd1-Halo with the trajectories of HaloTag-tagged constructs with known stoichiometries. We fused either one or two copies of the HaloTag to Sec61b, which is known to predominantly exist as an ER-resident monomer (Figure 4B). Thus, these constructs provide ground truth data for a perfect HaloTag monomer and dimer. We imaged U-2 OS expressing these constructs under the same conditions used for dcSMT of Hrd1-Halo. The extracted trajectories were algorithmically classified regarding correlation between both channels by analyzing prolonged proximity of detected molecules (Figure S3). While also directly visually apparent (Figure 4C, <u>Video S6</u>), this quantification highlighted the stark difference between the constructs. Whereas in Halo-Sec61b, only 1.6 ± 2.7 % of all trajectories were identified as correlated, marking the false-positive rate of the correlation classifier used, 17.4 ± 11.8 % of 2xHalo-Sec61b and 51.1 ± 13.7 % of Hrd1-Halo trajectories were classified as correlated (Figure 4D).

We calculated the expected relative amounts of correlated tracks for a hypothetical homo-tri- and homo-tetramer based on the labeling parameters extracted from the Halo-Sec61b and 2xHalo-Sec61b datasets (see Methods). This yielded predicted autocorrelations of 31.7% for tri- and 43.5% for tetrameric assemblies. The regular occurrence of dual-color correlation observed for Hrd1-Halo, to our knowledge unprecedented in dual-color tracking of single complexes, clearly demonstrates that it is assembled into complexes containing more than two copies of itself. Due to this stochastic analysis approach, our data could either be explained by a tetrameric assembly or a mixture of higher-order assemblies. However, the absence of any bleaching traces indicative of more than four copies favors a model of predominantly tetrameric complexes.

### Oligomerization is an Intrinsic Property of the Hrd1 Protein

The ability of dcSMT to capture and quantify higher-order oligomeric states of Hrd1 promises a novel perspective on the modular nature of the Hrd1 complex reported in literature. For example, previous work suggested that the Hrd1 interaction partner Fam8A1 governs the assembly of Hrd1 oligomers and thereby mimics the function of yeast Usa1.^21^ We employed siRNA-mediated silencing to reduce the cellular levels of Sel1L, Fam8A, and HERPUD-1 and -2, established components of the Hrd1 ERAD complex. The resulting effects on complex stoichiometry were directly observed by autocorrelation of Hrd1 using dcSMT (Figure 5A). Despite efficient downregulation of other Hrd1 complex components (Figure S4A), no significant change in autocorrelation was observed. Remarkably, the loss of these selected subunits of the Hrd1 ligase complex did not affect the formation of Hrd1-Halo homo-oligomers. Downregulation of Hrd1 itself was inefficient for reasons that remain unclear, as quantified by both western blot and single-molecule counting (Figure S4B) but did show slight decrease in Hrd1-Halo autocorrelation.

**Figure 5:**
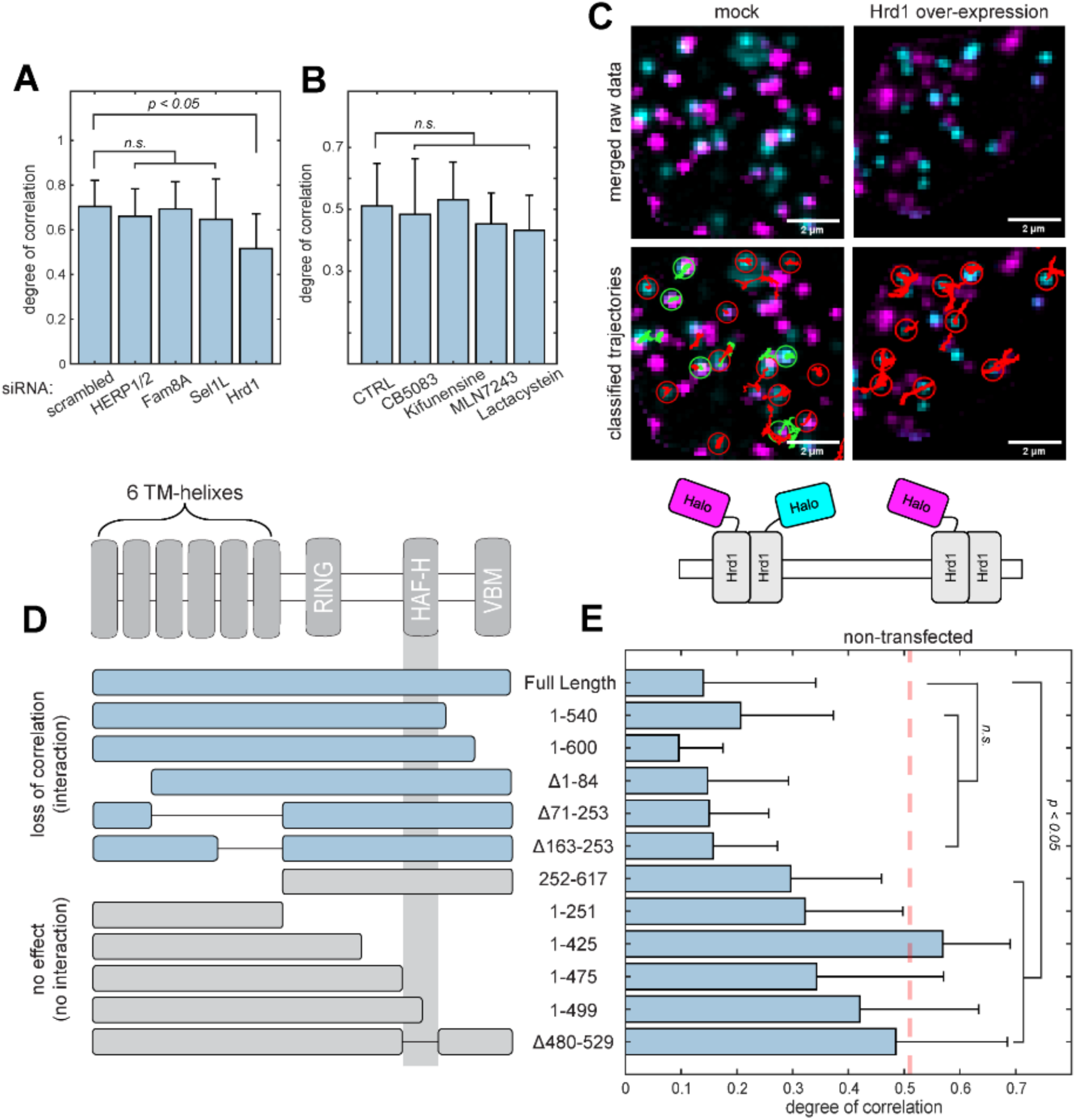
Competitive Overexpression and dcSMT shows that Hrd1 oligomerization is driven by Hrd1 480-529. **A)** Degree of cell-wise mean Hrd1-Halo autocorrelation in dcSMT after downregulation of defined Hrd1-complex components by 48h of siRNA treatment. Partial downregulation of Hrd1 leads to moderate but significant decrease in autocorrelation. A one-sided ANOVA was used to test statistical significance. n(cells) = 127/60/55/46/52 n(trajectories) = 8119/3603/2968/2895/2182. **B)** Degree of cell-wise mean Hrd1-Halo autocorrelation in dcSMT after prolonged inhibition (6h) of defined steps in ERAD. A one-sided ANOVA was used to test statistical significance. n(cells) = 66/36/38/20/23, n(trajectories) = 1582/643/958/416/495. **C)** Representative dcSMT recordings of Hrd1-Halo showing the loss of Hrd1-Halo autocorrelation upon plasmid-based overexpression of unlabeled Hrd1. The trajectories were color coded regarding the correlation between both channels (green = trajectory with correlation partner, red = uncorrelated). **D)** Schematic representations of Hrd1 truncations used for binding competition. **E)** Cell-wise mean Hrd1-Halo autocorrelation in dcSMT after plasmid-based overexpression of the indicated S-peptide tagged Hrd1 truncations. The red line marks the value for non-transfected cells. A one-sided ANOVA was used to test statistical significance. n(cells) = 29/15/13/18/24/18/33/14/8/18/22/15; n(trajectories)= 839/275/320/1690/1404/923/1389/1099/467/1001/1174/240.

Literature has also demonstrated changes in Hrd1 complex composition in response to several cellular cues, such as activation of the unfolded protein response (a cellular stress response program activated upon ER stress) and the targeted inhibition of ERAD.^20,36,37^ We revisited these observations with the capacity to directly observe complex stoichiometry. Surprisingly, treatment of Hrd1-Halo cells with inhibitors targeting either the proteasome (Lactacystin), ER mannosidase I (Kifunensine), VCP (CB5083) or ubiquitin-E1-ligase activity (MLN7243) for prolonged times (6 h) had no measurable effect Hrd1-Halo autocorrelation (Figure 5B). Moreover, overexpressing CD3-δΔ-HA, a Hrd1-specific substrate, did not result in significant changes of the fraction of correlated trajectories compared to control (Figure S5A). In summary, none of the previously described modulators of the ERAD-pathway cause any detectable change of Hrd1 homo-oligomerization.

Since we were not able to observe changes in Hrd1-Halo autocorrelation by modifying composition and function, we hypothesized that homo-oligomerization of Hrd1 may be an intrinsic property of the protein itself. We thus set out to develop a novel dcSMT-based assay capable of identifying protein interaction interfaces in situ. We hypothesized that overexpression of unlabeled Hrd1 would outcompete the endogenous Hrd1-Halo:Hrd1-Halo interaction and thus result in a reduction of Hrd1-Halo autocorrelation. Overexpression of truncated variants may thereby allow identifying regions within Hrd1 mediating this interaction, since variants lacking relevant stretches should be unable to outcompete the endogenous interaction. Thus, cells expressing these constructs should display a similar Hrd1-Halo autocorrelation as untreated cells.

Indeed, overexpression of S-peptide tagged full length Hrd1 in Hrd1-Halo cells led to a dramatic reduction in correlation in dcSMT (Figure 5C, <u>Video S7</u>), suggesting a dynamic component exchange between complexes exists that is too slow to be captured with SMT. We repeated these experiments using Hrd1-truncations missing several domains including pairwise deletions of the transmembrane helices and various truncations of Hrd1s cytosolic portion (Figure 5D/E, S6). Surprisingly, all TM-helix deletion constructs were able to outcompete and disassemble the endogenous Hrd1-Halo multimers. However, truncations even partially missing Hrd1_480-529_, a short, conserved peptide stretch in the cytosolic domain previously named the HAF-H domain, were unable to reduce the Hrd1-Halo dual-color correlation. Therefore, Hrd1 homo-oligomerization appears to be mainly driven by a short peptide within its cytosolic domain.

### Cytosolic Hrd1_480-529_ Forms a Tetrameric Helix Bundle Driving Oligomerization

Given that the tetrameric state of Hrd1 appears to be driven by the HAF-H domain, we aimed to characterize this interaction from a structural perspective. We began by using the homology-based prediction algorithm AlphaFold to model different substructures of Hrd1. ^38^ Monomeric Hrd1 was predicted to be composed of 8 bundled transmembrane helices and predominantly a disordered cytosolic domain. Within this disordered domain, AlphaFold predicted both the highly conserved ubiquitin-conjugating RING domain structure and an alpha helix formed by Hrd1_480-529_ (Figure S7A). Attempts to predict the structure of full-length Hrd1 multimers yielded mostly nonsensical results (Data not shown). Only dimeric Hrd1 led to a prediction with sufficient certainty of its TM-helices, most likely due to its close homology to the previously published yeast Hrd1 dimer. According to our competitive dcSMT assay, Hrd1_480-529_ driving Hrd1 oligomerization should reduce the complexity of the modeling, leading to more refined structural predictions for higher-order oligomers. We used AlphaFold to predict the structure of various multimers of Hrd1_480-529_ copies. While one to four copies of Hrd1_480-529_ led to acceptable pLDDT scores (Figure S7B), five or more copies dramatically reduced the quality of the prediction.

Dimeric Hrd1_480-529_ was predicted to form an antiparallel helix dimer (Figure 6A). This interaction appears to be driven by a hydrophobic patch, while the opposing face was composed of mostly polar residues. Hydrophobic patches are common features in protein-protein interaction interfaces and entropically drive complex formation. (consistent with this) In the predicted tetramer, the tetrameric helix bundle was composed of two helix dimers, each near-identical to the predicted dimer, interacted via their hydrophobic patches forming a hydrophobic core and a mostly polar surface. Aligning the amino acid sequence with the structural prediction revealed that Hrd_480-529_ contains various *heptad-repeats* (Figure 6B). This conserved amino acid motif is characterized by a repeating seven-residue sequence with defined positions for nonpolar amino acid side chains and is canonically involved in the formation of coiled-coil helix bundles.^39^ Interestingly, the hydrophobic core is flanked by two Arg503 and Tyr524 pairs in sufficient proximity to form hydrogen bonds between opposing helix pairs (Figure 6C). Arg503 was previously suggested to be involved in the interaction with Fam8A related to Hrd1s oligomerization.^21^

**Figure 6:**
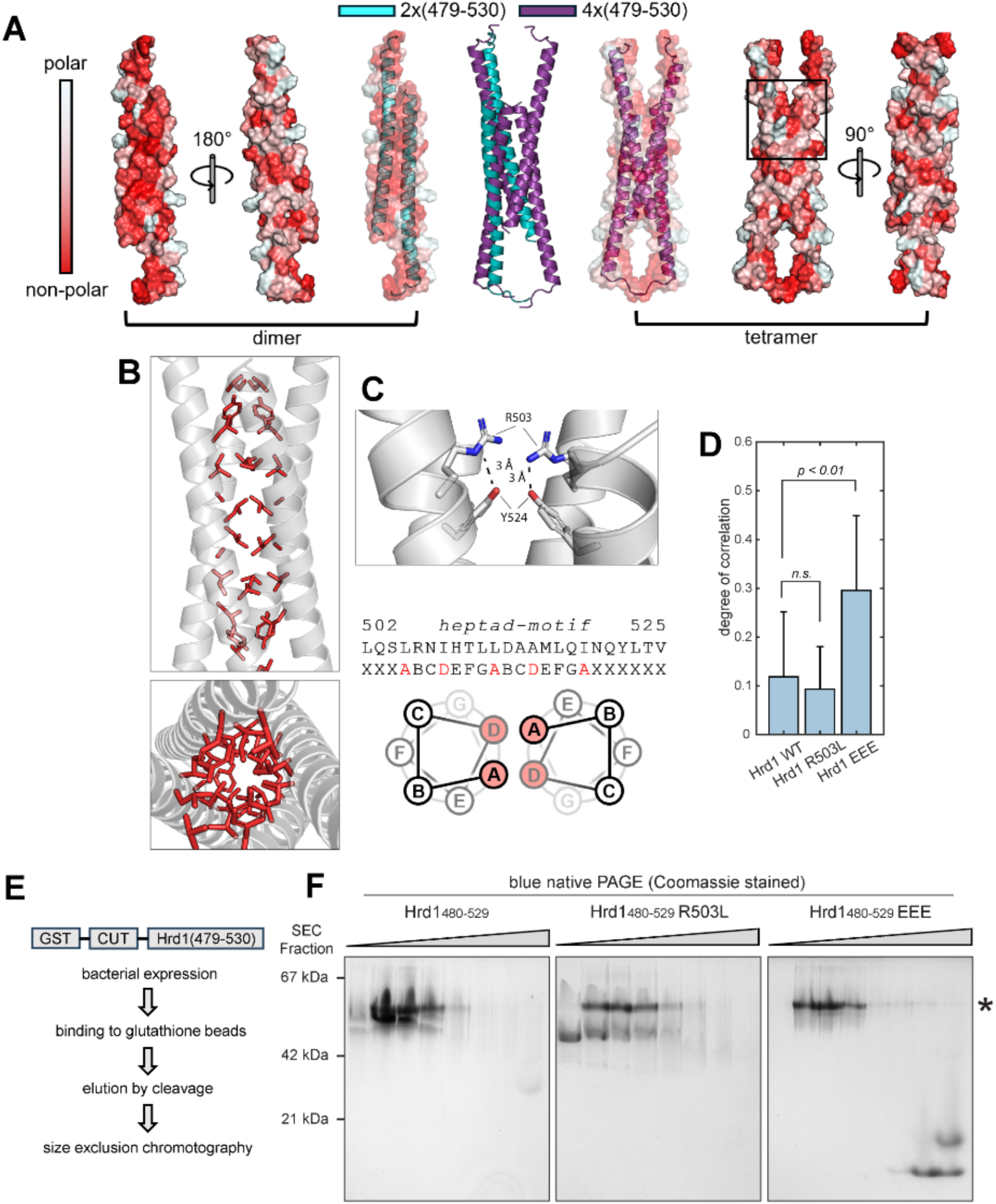
Hrd1 tetramerization is driven by a canonical coiled-coil helix bundle formed by Hrd1 479-530. **A)** AlphaFold predictions of dimeric and tetrameric Hrd1 479-530. The surface representation was color coded according to the hydrophobicity of the residue sidechains. **B)** Different representations of the hydrophobic core identified in the tetramer prediction. Red side chains mark amino acids partaking in the canonical heptad-repeats that form the hydrophobic cluster. **C)** Potential electrostatic interaction between R503 and Y524 observed in the tetramer prediction. **D)** Mean Hrd1-Halo autocorrelation in dcSMT after plasmid-based overexpression of the indicated constructs. In Hrd1 EEE three of the hydrophobic amino acids forming the postulated hydrophobic core were replaced by glutamic acid. A one-sided ANOVA was used to test statistical significance. n(cells) = 26,26,31 n(trajectories) = 872,917,1173. Workflow used for recombinant expression and purification of Hrd1 479-530 from *E. coli*. **E)** Coomassie-stained blue native PAGE of purified Hrd1 479-530 variants after size exclusion chromatography. The Asterisk indicates residual cleaved GST-tag present in all samples.

To prove the relevance of the predicted tetrameric helix bundle for Hrd1 oligomerization in cells, we aimed to disrupt the proposed hydrophobic core by targeted mutagenesis. Using dcSMT, we monitored its effect on Hrd1 oligomerization in our previously described competitive interface mapping approach. Based on the AlphaFold prediction, we generated two Hrd1 variants that we predicted to be compromised in their ability to form the helix bundle: in Hrd1 L514E I521E L528E (Hrd1 EEE), three of the nonpolar residues in the *heptad-repeats* were mutated to charged glutamic acids to disrupt the hydrophobic core. In Hrd1 R503L, the potential hydrogen bond between Arg503 and Tyr524, previously reported to be involved in Hrd1s interaction with Fam8A1, was disrupted. We probed the effect of these substitutions on the constructs’ propensity to outcompete the endogenous Hrd1-Halo interaction in dcSMT (Figure 6D). While Hrd1 R503L led to a near-complete abolishment of correlation similarly as Hrd1 FL, Hrd1 EEE had little to no effect on the Hrd1-Halo correlation and was similar in its effect to overexpressed Hrd1 Δ480-529.

While this clearly showed that these polar residues are central for the establishment of Hrd1 multimers, it only indirectly confirmed the four-helix bundle as predicted by AlphaFold. To obtain more direct evidence, we aimed to characterize this motif using classical biochemistry. We recombinantly expressed and purified Hrd1_480-529_, Hrd1_480-529_ R503L, and Hrd1_480-529_ EEE from *E. coli* and analyzed their tendency for oligomerization using a combination of MALDI/TOF MS and blue native- and SDS-PAGE. GST-tagged Hrd_480-529_ was isolated from lysate by immobilization on glutathione beads and eluted by addition of GST-tagged PreScission protease (Figure 6E). The resulting eluates were separated using size-exclusion chromatography (SEC) and analyzed using native PAGE (Figure 6F, Figure S8). The SEC inputs of both samples were analyzed using MALDI/TOF MS (Figure S9), confirming that the sample exclusively contained cleaved GST-tag, that could not be removed during the purification, and the corresponding Hrd1 peptide. While Hrd1_480-529_ and Hrd1_480-529_ R503L behaved very similarly, Hrd1_480-529_ EEE showed fundamental differences in native PAGE and SEC: while Hrd1_480-529_ EEE eluted in late SEC fractions and migrated with an apparent molecular weight of 12 kDa in native PAGE, Hrd1_480-529_ eluted earlier and migrated as a 50 kDa species, migrating just below the dimeric 56 kDa GST-tag. When the samples were separated on denaturing SDS-PAGE, the high molecular weight band of Hrd1_480-529_ instead migrated mainly as a species equivalent to monomeric Hrd1_480-529_ EEE and a faint second band equivalent to an assembly of twice the molecular weight. Taken together, purified Hrd1_480-529_ assembled in highly stable, partially SDS-resistant tetramers. This tetrameric assembly was dependent on the nonpolar side chains driving the formation of the predicted coiled-coil helix bundle. Replacing them with polar residues as in Hrd1_480-529_ EEE resulted in a behavior as expected for the monomeric peptide. Consistent with its ability to outcompete the endogenous Hrd1-Halo oligomerization, purified Hrd1_480-529_ R503L was indistinguishable from Hrd1_480-529._ Therefore, this biochemical characterization was in line with both the quantitative dcSMT and the structural prediction, demonstrating Hrd1s assembly in predominantly stable homo-tetramers.

## Discussion

### Hrd1 is Assembled in Stable Homo-tetramers

In this work, we combine dcSMT, structural predictions, and in vitro biochemical analysis to describe and characterize the tetrameric nature of the Hrd1 complex. We identify an amphipathic helix on the cytosolic face of Hrd1 that is responsible for the formation of higher-order homo-oligomers in situ and a tetrameric complex in vitro. The cytosolic domain of Hrd1 has been well established in conveying roles beyond ubiquitination using both functional assays and immunoprecipitations. ^21,40^ However, the structures resolved for Hrd1 are limited to its transmembrane domains since the disordered cytosolic domain interferes with purification procedures. Thus, a mechanistic understanding of its contribution to the holo-complex and Hrd1 function is missing. The SMT approaches presented here permit a novel, unbiased perspective on the stoichiometry of the full-length holo-complex, which has been hidden from classical biochemistry and structural biology due to their dependence on purified and in vitro systems. In this work, we present four independent lines of evidence demonstrating mammalian Hrd1s assembly in homo-tetramers: (1) the measured degree of correlation of Hrd1-Halo in dcSMT exceeds the measured correlation of a dimeric construct and the theoretical degree of correlation of a trimer, (2) events bleaching in 3 or more steps are regularly observed, (3) AlphaFold predicts a tetrameric assembly of Hrd1_480-529_ with high accuracy, and (4) purified Hrd1_480-529_ migrates as a homo-tetramer in native PAGE. This homo-tetramerization is an intrinsic property of Hrd1 and independent of other complex components or ERAD activity, since neither siRNA-mediated targeted knockdown nor specific ERAD inhibitions had any effect on Hrd1 oligomerization as measured by dcSMT.

### Implications of Higher-order Hrd1 Assembly

The relevance of higher-order oligomers of Hrd1 in protein dislocation has been previously suggested based on high molecular weight complexes observed in in vitro experiments, but the idea lost traction due to the solved structures of mono- and dimeric yeast Hrd1. However, Derlin-1 has been recently described as being assembled into homo-tetramers,^15^ and isolated yeast Hrd1 complexes have been observed to be composed of three and more copies by in vitro single-molecule imaging.^41^ Thus, our data supports a line of evidence in the literature that the functional form of the Hrd1 complex is a higher-order structure.

The observed tetrameric nature of the Hrd1 complex suggests interesting possibilities for the mechanism of Hrd1-mediated retro-translocation. In contrast to co-translational protein insertion, the dislocation process centered around Hrd1 faces a wide variety of substrates with different topologies and even dislocates partially folded and glycosylated substrates. Published Hrd1 complex structures are sterically unable to transport anything bigger than a single unfolded peptide. On the other hand, a high-molecular-weight assemblies of Hrd1 could harbor even bulky and modified substrates, such as the glycosylated forms that are inherent to most Hrd1 substrates. A tetrameric holo-complex composed of Hrd1-Sel1L-Derl1-Fam8A-HERPUD1 complex would encompass a total of (6+1+6+3+2)x4 = 72 transmembrane helices and a total molecular weight of (68+89+28+44+43)x4 = 1088 kDa. A complex of this size composed of dynamically interacting transmembrane domains may facilitate more sterically demanding dislocation mechanisms that would not require complete unfolding of the substrate and permit passage of the glycosylated form.

### Role of Hrd1s cytosolic domain in complex assembly

We show that Hrd1s homo-oligomerization is driven by a peptide residing in its cytosolic domain forming a highly stable intermolecular coiled-coil helix bundle. Interestingly, the Hrd1 complex is known to utilize a different subset of interaction partners depending on the substrate, suggesting a dynamic assembly of the complex. Consistent with this, electrophysiological measurements suggested a flexible pore size for Hrd1.^37^ The location of its highest affinity interaction at the end of a vast disordered region offers an intriguing explanation for the modularity of the Hrd1 complex. Hrd1 tightly interacting via its cytosolic domain, while ensuring the high local concentrations required for its function, could allow transient interaction and dynamic rearrangement of its membrane segments dependent on a substrate’s requirements. This dynamic architecture could also explain the heterogeneity of Hrd1 complexes when probed biochemically.

In contrast to yeast, where Hrd1 oligomerization is mediated by Usa1,^19^ mammalian Hrd1 oligomerization is an intrinsic property. Interestingly and accordingly, Usa1 is the only yeast Hrd1 complex component that appears to not have a direct mammalian homolog. We propose that the Hrd1-tetramer forms a stable central scaffold for the formation of the holo-complex, and the binding of other complex components relies on its homo-oligomerization. Mammalian Hrd1 variants missing HAF-H are defective in homo-oligomerization, substrate dislocation, and interaction with Fam8A1 or HERPUD1.^21^ Similar observations have been made in yeast: Usa1-dependent Hrd1 oligomerization is required for its interaction with Derl1. Thus, in mammals, the HAF-H domain may functionally replace the role of yeast Usa1.

Due to the dynamic nature of ERAD, understanding retro-translocation has been a challenge for decades. Here, we demonstrate that single-molecule-based imaging can provide structural insights that overcome key limitations in classical biochemistry, offering previously unachievable perspectives on Hrd1-mediated dislocation and dynamic protein complexes in general.

## Methods

### Cell culture

U-2 OS cells were cultured in DMEM containing 10% v/v fetal bovine serum (FBS) supplemented with penicillin and streptomycin incubated at 37°C and 5% CO_2_. Cells were supplied by ATTC and regularly tested for mycoplasma infections using a PCR-based kit (Thermo Fisher).

### Generation of Monoclonal Cell Lines using CRISPR-Cas9

Monoclonal CRISPR/Cas9 edited cell lines were generated following the protocol provided in Ran et al. 2013 using a single-plasmid CRISPR/Cas9 system^5^ followed by fluorescence-activated cell sorting (FACS) and single cell expansion. For HaloTag knockin, a repair template coding for the HaloTag flanked by 1000-bp arms targeting the Hrd1 locus was co-transfected. U-2 OS cells were plated in T25 cell culture flasks (Sarstedt). At 24 hours post-transfection, cells were trypsinized, pelleted (300 rcf, 5 min), and resuspended in PBS. Cell clumps were removed by filtration through a round-bottom polystyrene test tube with cell strainer (Fisher Scientific) and kept on ice until FACS. Using a BD FACSAria™ III Cell Sorter (Biosciences), single mCherry-positive cells (610/20 filter) were sorted into 96-well plates (Sarstedt) containing 100 µl DMEM with 10% FCS and 50 µg/ml gentamicin (Sigma-Aldrich). Single clones were expanded into larger culture formats as they reached confluency. Editing was confirmed by immunoblotting for Hrd1 and targeted sequencing.

### Plasmid and siRNA Transfection

Plasmid DNA was transfected into U-2 OS cells using GenJet^TM^ (SignaGen) 24 h post-seeding according to the manufacturers’ protocol. For 35-mm dishes 1 µg DNA and 3 µl GenJet^TM^ was used; for T25 flasks, 5 µg DNA and 15 µl GenJet^TM^ was used. For co-transfections, the total DNA amount was kept constant and plasmids were mixed 1:1. ON-TARGETplus siRNA Smartpool (Horizon Discovery) transfection was performed using DharmaFECT^TM^ 2 (Horizon Discovery) following the manufacturers’ protocol with a final concentration of 50 µM siRNA and 2 µl DharmaFECT^TM^ 2 for 35-mm dishes. Samples were collected or imaged 48 h post-transfection.

### Lentivirus Preparation and Transduction

Emerald-Sec61b and the Halo-Sec61 reference constructs were delivered by lentiviral transduction. Lentivirus was produced in HEK293T cells by seeding 5 × 15 cm dishes with 1.1 × 10^7^ cells one day prior to transfection. Per dish, 3.5 µg VSVG, 2.5 µg Rev, 5 µg Pol, and 12 µg pLJM1 were mixed in 450 µl D10, followed by addition of 36 µl PEI. The mixture was incubated for 20 min at RT and added to the cells. The next day, the medium was replaced with 15 ml fresh D10. Medium was collected and replaced with fresh 15 mL D10 at 8, 12, and 20 h. Collected media were centrifuged at 500 × g for 3 min and filtered through a 0.22 µm Millex^®^ syringe filter (Merck). Virus was pelleted by centrifugation at 47,850 × g for 2 hours at 4°C, resuspended in 500 µl D10, and aliquoted (50 µl) for storage using liquid nitrogen.

For transduction, U-2 OS cells were seeded as previously described. After 24 h, the medium was replaced with 1.35 ml D10 containing 1 µg/ml polybrene infection reagent (Sigma-Aldrich). After 1 h, 150 µl virus solution in D10 containing 1 µg/ml polybren was added. Viral titers were estimated using serial dilution, but typically a final virus dilution of 1:30 yielded appropriate transduction efficiency. Cells were transduced for 8 h before imaging.

### Radioactive pulse-chase assays

Degradation of model substrate CD3-δΔ-HA was followed using radioactive pulse-chase assays followed by immunoprecipitation (PC/IP). Cells were transfected 24 hours prior to the experiment. For radioactive labeling, cells were washed with PBS and incubated in starvation medium (DMEM w/o cysteine/methionine/glutamine + 5 mM L-glutamine + 10% FCS) for 10 min at 37°C. The medium was then replaced with labeling medium (starvation medium supplemented with 1.7 MBq/ml Expre^35^S^35^S Protein Labeling Mix (Perkin Elmer)) and incubated for 15 min at 37°C. Cells were rinsed twice with PBS and incubated with chase medium (D10 + 48 mg/L L-Cystein + 30 mg/L L-methionine).

Samples were taken at listed time points by rinsing with PBS and adding 300 µl lysis buffer. Lysates were rotated at 4°C for 1 h, followed by centrifugation (15,000 × g, 4°C, 20 min) to remove residual debris. For immunoprecipitation, 1.2 ml lysis buffer, 15 µl Protein A–Sepharose^TM^ 4B (Thermo Fisher), and 1 µl anti-HA antibody were added, and the samples were incubated overnight at 4°C. The next day, agarose beads were pelleted (500 × g, 30 s) and washed three times with lysis buffer. Bound proteins were eluted from the pelleted beads by addition of 25 µl 1× SDS sample buffer, separated by SDS-PAGE, and the resulting gel was fixed in 10% (v/v) acetic acid, washed with ddH_2_O, and dried on Whatman paper using a Model 583 gel dryer (Bio-Rad Laboratories, Inc.). Dried gels were exposed to a BAS Storage Phosphor Screen (GE Healthcare) for 48-96 h, signals were detected using a Typhoon FLA 9500 laser scanner (GE Healthcare), and band intensities were analyzed using the manufacturer’s software.

### SDS-PAGE and Western Blotting

Protein samples from cell lysis, immunoprecipitation, or protein purification were analyzed using sodium dodecyl sulfate-polyacrylamide gel electrophoresis (SDS-PAGE). Proteins were separated by molecular weight using a gel electrophoresis system (Hoefer Inc.) filled with Laemmli running buffer (LRB). Proteins were separated by 3% stacking gel (125 mM Tris-HCl (pH 6.8), 3% (v/v) acrylamide, 0.15% (v/v) bisacrylamide, 0.1% (w/v) SDS, 0.25% (v/v) TEMED, and 2.5% (w/v) APS) followed by 12% or 18% separating gel (500 mM Tris-HCl (pH 8.8), 12% or 18% (v/v) acrylamide, 0.006% or 0.009% (v/v) bisacrylamide, 0.1% (w/v) SDS, 0.25% (v/v) TEMED, and 2.5% (w/v) APS). Electrophoresis was performed at 80 V until the dye front reached the separating gel, then 120 V for 90 min.

For immunoblotting, proteins were transferred from SDS-PAGE gels onto methanol-activated PVDF membranes using a TE22 Mighty Small Transfer Tank (Hoefer Inc.) containing transfer buffer and applying an electric current of 250 mA for 90 minutes, then sandwiched between Whatman paper. Membranes were blocked in 10% (w/v) skim milk powder in TBST for 30 min at RT to reduce nonspecific antibody binding. Primary antibody solutions were prepared in 5% skim milk powder in TBST and applied to the blocked membranes at 4°C overnight. The membranes were then washed three times using 10 ml TBST and three times using 10 ml PBS (5 min each). HRP-conjugated secondary antibodies were freshly prepared and applied to the membranes for 1 hour at RT followed by washing as previously described. Signal detection was performed using *Western Lightning® Plus-ECL, Enhanced Chemiluminescence Substrate* (PerkinElmer, Inc.) and imaged using an Odysee® Fc imaging system (LI-COR Biosciences).

### Co-Immunoprecipitation

1×10^7^ U2-OS and U2-OS Hrd1 HaloTag cells were lysed in 1 ml ice cold RIPA buffer supplemented with protease inhibitor (Roche, complete ULTRA protease inhibitor tablet, 1 tablet per 10 mL) and Benzonase on ice. Lysates were rested on ice for 30 min. The protein concentration was quantified using Pierce Bicinchoninic Acid Protein Kit (Thermo Fisher Scientific) and normalised across conditions. 500 μg of protein was loaded onto pre-equilibrated bead slurry (TrueBlot Anti-Rabbit Ig IP Agarose Beads, #00-8800-25) and incubated on a rocking platform for 60 minutes at 4°C for preclearing. 5 μg anti-Hrd1 antibody (Proteintech, #13473-1-AP) was conjugated to Rockland TrueBlot Agarose beads per sample, then protein lysate from the preclearing was loaded and immunoprecipitated overnight at 4°C. The beads were then washed three times with IP wash buffer (50 mM NaCl, 10 mM Tris, pH 7.5) at 4°C. Bound proteins were eluted from the beads via heating at 65°C for 10 min and vortexing in elution buffer (RIPA buffer supplemented with 4% SDS and 1× Laemmli buffer), before being subjected to western blot.

### Recombinant production of human Synv1 proteins and oligomerization analysis

Constructs for wild-type and variant (R503L, L514E/I521E/L528E) SYVN1 comprised amino acids 487-529 of human Syvn1 (UniProt: Q86TM6). Genes were cloned via restriction into the pGEX-6P1 vector, producing N-terminal GST-tagged fusion proteins with a PreScission protease cleavage site. Proteins were expressed using *E. coli* T7 Express cells (NEB) co-transformed with pRARE2. Cultures were grown in TB medium supplemented with 100 µg/mL ampicillin and 34 µg/mL chloramphenicol at 37°C using the LEX (“Large-scale EXpression”) bioreactor technology (Epiphyte3 Inc., Toronto, Canada) until OD600 reached ∼2.5. Gene expression was induced with 0.5 mM isopropyl β-D-1-thiogalactopyranoside at 17°C and cultures were grown overnight at 17°C. Cells were harvested by centrifugation and pellets were stored at -70°C.

For purification, cells were resuspended in lysis buffer (50 mM Tris pH 7.5, 0.15 M NaCl) supplemented with 0.25% (w/v) 3-[(3-cholamidopropyl)-dimethylammonio]-1-propane-sulfonate, 1 mM phenylmethyl-sulfonyl fluoride, 0.5 mM dithiothreitol (DTT), 8240 U/mL lysozyme, and 17 U/mL benzonase. Cells were lysed by repeated freeze-thaw cycles, and lysates were clarified by centrifugation. GST-fusion proteins were captured from the supernatant by incubation with glutathione sepharose 4B affinity chromatography resin (Cytiva) equilibrated with lysis buffer for 1.5 h (at 4°C?). Protein-bound resin was washed once with lysis buffer and three times with lysis buffer with 1 M NaCl to remove contaminating proteins and nucleic acids. Next, protein-bound resin was re-equilibrated in lysis buffer with 1 mM DTT and GST-tagged PreScission protease and incubated overnight at 8°C to facilitate GST fusion tag cleavage and elution of target proteins. Purified proteins were analyzed by semi-analytical gel filtration chromatography on a Superdex 200 Increase 10/300 GL column (Cytiva) equilibrated with 20 mM Hepes-NaOH pH 7.5 and 0.2 M NaCl. Eluted fractions were further analyzed by blue native (BN)-PAGE using 20% Bis-Tris gels and SERVAGel N Native Gel Kit (Serva), following the manufacturer’s protocol.

### Coomassie Staining

Proteins were visualized by staining gels with Coomassie staining solution (10 % (v/v) acetic acid, 40 % (v/v) methanol, 0.25 % (w/v) Coomassie Brilliant Blue R-250) for 30 min at RT. Gels were destained in 10 % (v/v) acetic acid, 40 % (v/v) methanol until protein bands were clearly visible. Images were acquired using the Odyssey® Fc imaging system (LI-COR Biosciences).

### Coverslip Preparation and Coating

Glass coverslips (35 mm, thickness 1.5; Roth) were plasma-cleaned for 10 min at 40% power using room air. The plasma-facing side was tracked throughout the cleaning process. Coverslips were submerged in 5% (v/v) HELLMANEX III (Hellma Analytics) in ddH_2_O and sonicated for 5 h, rinsed three times with ddH_2_O, submerged in ddH_2_O, sonicated for an additional 5 h, rinsed again three times with ddH_2_O, submerged in ethanol, and air-dried in a sterile hood. Dried coverslips were coated by applying 200 µl 10% (v/v) Matrigel (Corning) in PBS for 30 min, rinsed with 10 ml PBS, and stored in PBS in 35 mm dishes until cell seeding.

### Spinning Disk Imaging

Spinning disk imaging was performed using a CSU-W1 Spinning Disk Unit attached to an inverted microscope stand CSU-W1 with a 50 µm pinhole and a Borealis unit (Yokogawa). Illumination was performed by a laserbox equipped with 488 nm (150 mW), 561 nm (100 mW) and 637 nm (140 mW) lasers using a dichroic beamsplitter 405/488/561/640 (Chroma Technology). Images were acquired using 40× oil immersion (CFI P-Fluor/NA1.3/WD 0.2) and 100× oil immersion (CFI P-Apo λ /NA 1.45/WD 0.13) objectives. Temperature and CO_2_ were maintained by an OkoLab stage incubator. Emission was collected with an EMCCD Andor iXON DU-888 and filtered through a quad-band emission filter 446/523/600/677 (Chroma Technology). Exposure time (100-500 ms) and laser intensities were individually adjusted: 1-2% (488 nm, mEmerald-Sec61b) and 20-40% (647 nm, Hrd1-Halo).

### Total Internal Reflection (TIRF) and SMT

TIRF imaging was performed with a Nikon Eclipse Ti2-E inverted research microscope. The microscope operated with a CFI HP Plan Apochromat TIRF 100xAC/1.49 NA oil objective and a LU-NC-E laser unit composed of 488 nm (70 mW), 561 nm (70 mW), and 640 nm (125 mW) lasers using a TI-LA-TIRF module for laser inclination. Bandpass filters (Chroma Technology) used included: F47-525 (525/50 nm), F47-595 (595/50 nm), and F47-700 (700/75 nm). Four Andor iXON – DU 897U EMCCD cameras (512 × 512 with 16 µm pixel) were used for detection. Signals were separated using a three MultiCam camera splitter with dichroic mirrors (AHF Analysetechnik): F48-486, F48-559, F48-644. Imaging of Hrd1-Halo was performed with an exposure time of 20 ms only imaging the central 128 × 128 field of view. Laser intensities were set to 1% (488 nm), 70% (549 nm), and 70% (647 nm).

### HaloTag Labeling

For fluorescent labeling, HaloTag-expressing cells were plated in 35-mm dishes, rinsed with 10 ml PBS, and incubated with 1 mL antibiotic free D10 with 250 nM Halo-JF549 and/or 1 µM Halo-JF646 for 10 min. After labeling, cells were rinsed with 30 mL PBS, incubated in 2 mL D10 for 20 min, rinsed again with 10 mL PBS, and stored in phenol red-free D10 until imaging.

### Single-particle Localization and Tracking

Localization and tracking were performed using the ImageJ plugin Trackmate ^6,7^. For localization, the LoG detector was used with an expected object diameter of 750 nm and individually adjusted quality thresholds. Localizations were linked using a simple LAP tracker with an allowed distance and gap of 800 nm per step and two linking gaps allowed. Trajectories with less than 20 steps were excluded from all analysis.

### Dual-Color Correlation

Correlated movement in dual-color recordings was analyzed using self-written scripts in MATLAB after localization and tracking in Trackmate. For proximity correlation, the nearest neighbor in the 549-channel was found for every localization within each trajectory of the 647-channel. If the distance was below the manually set distance threshold, the localization was considered correlated. For directional correlation, the dot product was computed between every step in the 647-trajectories and all steps occurring in the 549-trajectories within 2 µm after vector normalization. If the highest dot product of a step in the 647-trajectories exceeded a manually set threshold, it was considered correlated. This procedure resulted in binary traces describing the 647-trajectories, which indicated if a step within this trajectory had a correlation partner or not. These traces were then analyzed regarding consecutive correlations. A trajectory was flagged as correlated if a sufficient number of consecutive steps were correlated. This was expanded by tolerating gaps in this sequence. Comparing these flags to a manually generated ground truth allowed calculation of the true negative rate, the true positive rate, and the false positive rate of the classifier and were used for optimization (Supplementary Figure 6). All shown analysis were performed using the proximity correlation approach using a maximal distance of 150 nm, 1 allowed gap, and proximity for 7 consecutive steps. From that, the degree of correlation was calculated as the fraction of correlated trajectories compared to all trajectories in the 647-channel.

#### Simulation of higher-degree oligomers

To predict the degree of correlation for a di-, tri-, or tetrameric assembly, a multinominal distribution with the states A (color 1), B (color 2), and U (unlabeled) was assumed. The fraction of correlated trajectories in color A was calculated as:

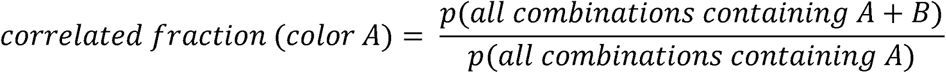

For a dimeric assembly (multinomial, two trials), this corresponds to:

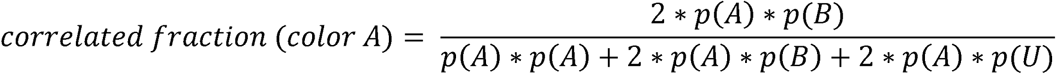

The labeling ratio *r* between A and B can be estimated based on the number of trajectories in the Halo-Sec61b dataset, which then results in:

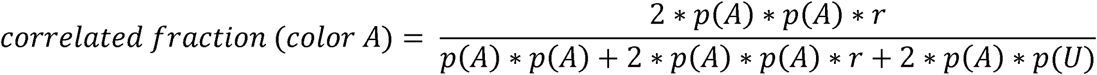

Assuming that

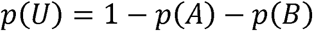

which equals

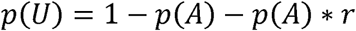

results in

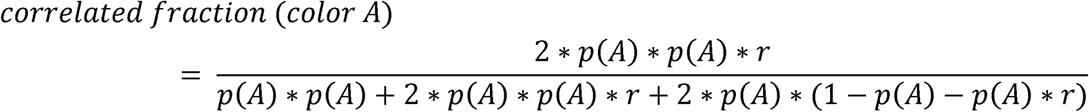

Since the correlated fraction of a dimer is known based on the analysis of the 2×Halo-Sec61b dataset, this can be used to calculate p(A) and therefore p(B) and p(U) given the labeling ratio *r*. The expected correlated fraction for higher order oligomers multinomial distribution with three or four trials can then be calculated as:

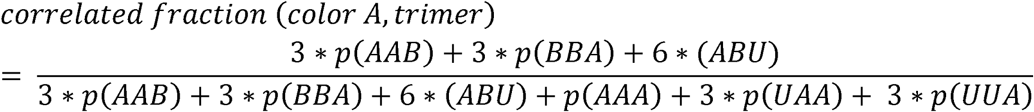

and

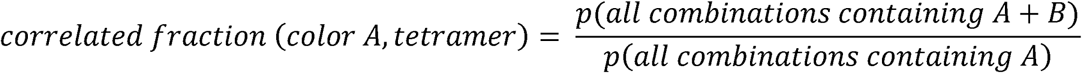

with

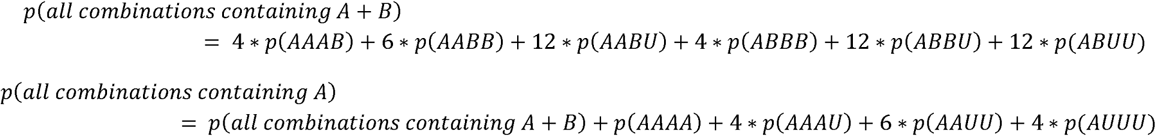

Inserting the values previously estimated based on Halo-Sec61b and 2×Halo-Sec61b tracking yields the theoretically expected degree of correlation.

### Structural prediction using AlphaFold

The structural predictions were performed using a publicly available implementation of AlphaFold2 (DeepMind) with the sequence FAGLTPEELRALEGHERQHLEARLQSLRNIHTLLDAAMLQINQYLTVLASLG using the default parameters. Images of structures were generated using PyMOL and the pLDDT values were plotted using MATLAB.

### Statistical testing and Plotting

Statistical tests were performed using MATLAB. Statistical significance was tested using the Tukey-Kramer pairwise comparison test, comparing each condition to the control condition implemented using a combination of MATLAB ANOVA and *multcompare*. For the degree of correlation, the mean cell-wise degree of correlation were compared, whereas in case of the diffusion coefficients, all diffusion coefficients from one dataset were pooled and compared. Plotting and calculations were performed using custom written scripts in MATLAB. All error bars shown in this work depict one standard deviation.

#### Plasmids

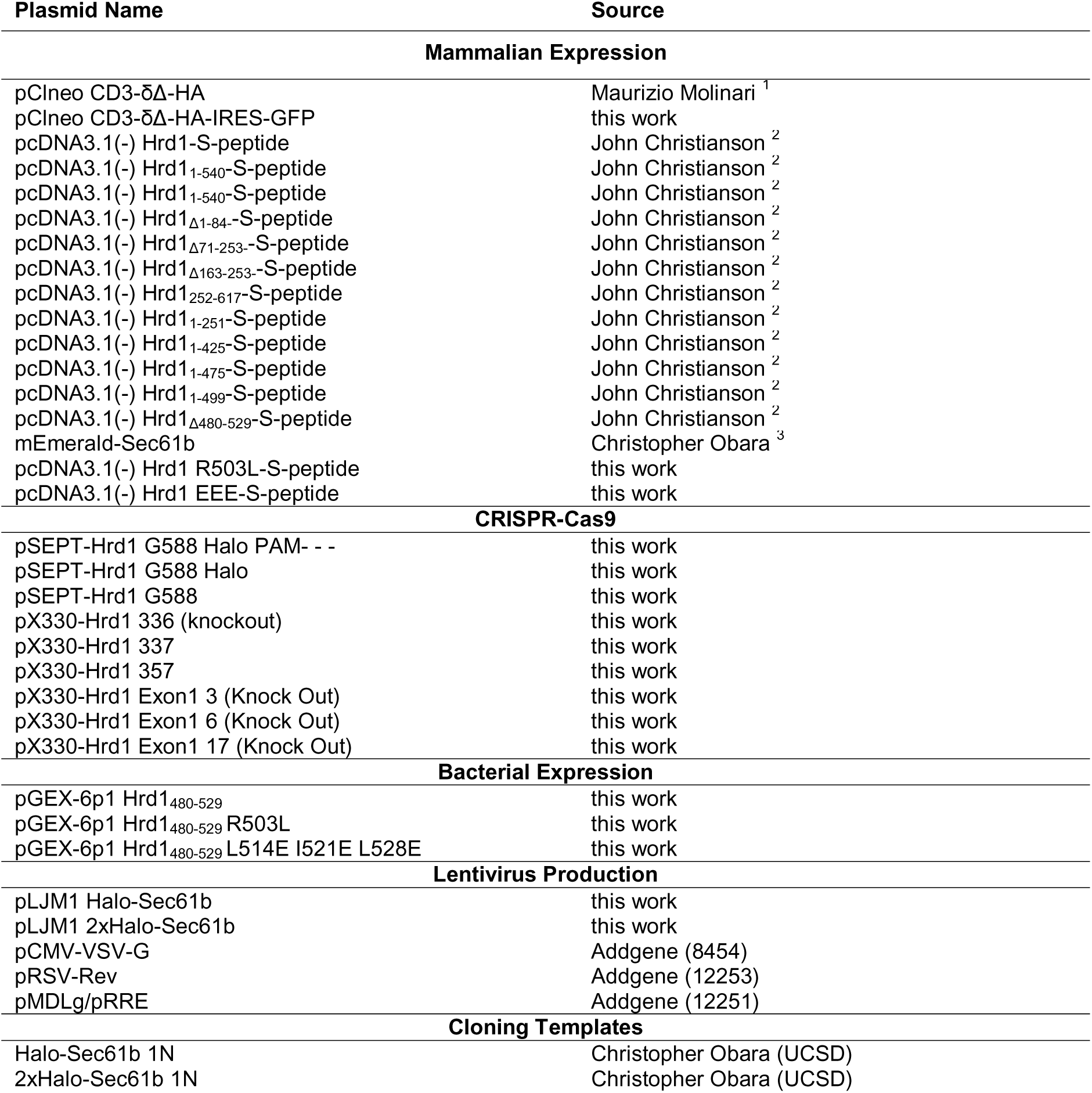
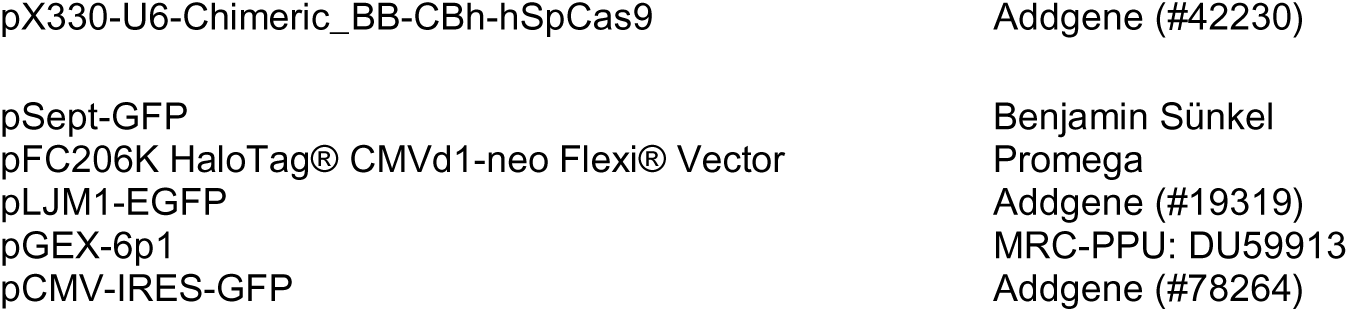

#### Antibodies

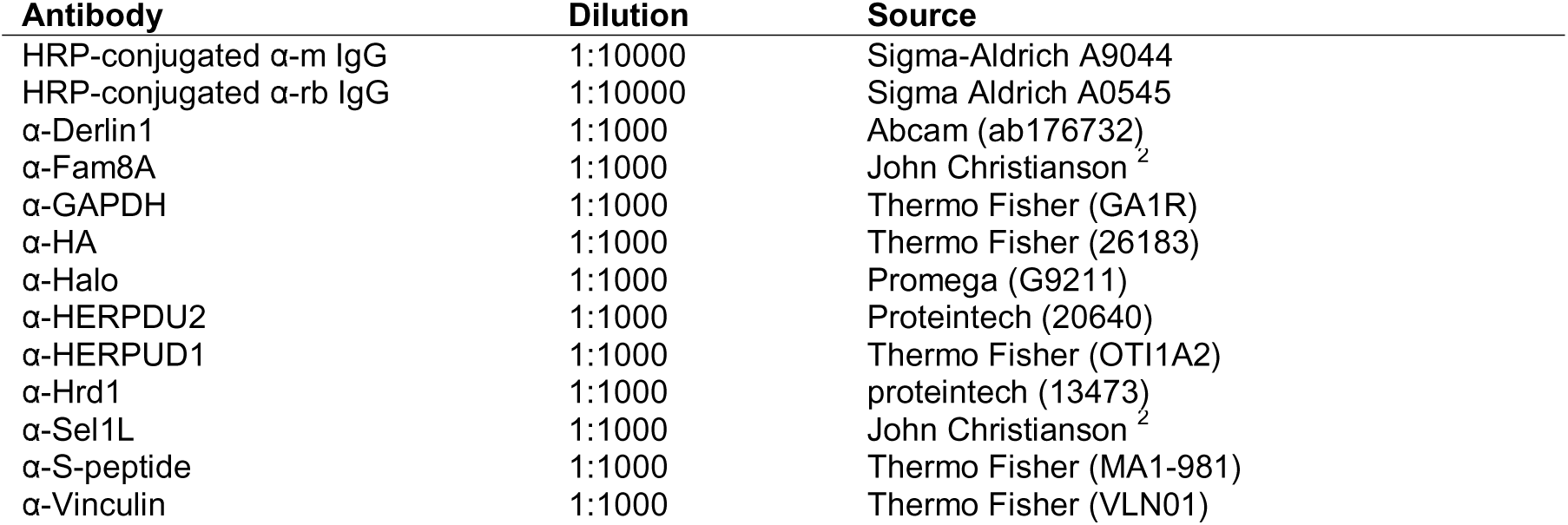

## Bibliography

1. Uhlén, M., Fagerberg, L., Hallström, B.M., Lindskog, C., Oksvold, P., Mardinoglu, A., Sivertsson, Å., Kampf, C., Sjöstedt, E., Asplund, A., et al. (2015). Tissue-based map of the human proteome. Science (1979) 347. 10.1126/science.1260419.

2. Sun, Z., and Brodsky, J.L. (2019). Protein quality control in the secretory pathway. Preprint at Rockefeller University Press, https://doi.org/10.1083/jcb.201906047 10.1083/jcb.201906047.

3. Christianson, J.C., and Carvalho, P. (2022). Order through destruction: how ER-associated protein degradation contributes to organelle homeostasis. EMBO J 41. 10.15252/embj.2021109845.

4. Meyer, H., Wang, Y., and Warren, G. (2002). Direct binding of ubiquitin conjugates by the mammalian p97 adaptor complexes, p47 and Ufd1-Npl4. EMBO J 21.

5. Fenech, E.J., Lari, F., Charles, P.D., Fischer, R., Laétitia-Thézénas, M., Bagola, K., Paton, A.W., Paton, J.C., Gyrd-Hansen, M., Kessler, B.M., et al. (2020). Interaction mapping of endoplasmic reticulum ubiquitin ligases identifies modulators of innate immune signalling. Elife 9, 1–29. 10.7554/eLife.57306.

6. Bernasconi, R., Galli, C., Calanca, V., Nakajima, T., and Molinari, M. (2010). Stringent requirement for HRD1, SEL1L, and OS-9/XTP3-B for disposal of ERAD-LS substrates. Journal of Cell Biology 188, 223–235. 10.1083/jcb.200910042.

7. Wei, X., Lu, Y., Lin, L.L., Zhang, C., Chen, X., Wang, S., Wu, S.A., Li, Z.J., Quan, Y., Sun, S., et al. (2024). Proteomic screens of SEL1L-HRD1 ER-associated degradation substrates reveal its role in glycosylphosphatidylinositol-anchored protein biogenesis. Nat Commun 15. 10.1038/s41467-024-44948-2.

8. Miao, X., Wu, J., Chen, H., and Lu, G. (2022). Comprehensive Analysis of the Structure and Function of Peptide:N-Glycanase 1 and Relationship with Congenital Disorder of Deglycosylation. Preprint at MDPI, https://doi.org/10.3390/nu14091690 10.3390/nu14091690.

9. Zhao, D., Wu, X., and Rapoport, T.A. (2025). Initiation of ERAD by the bifunctional complex of Mnl1/Htm1 mannosidase and protein disulfide isomerase. Nat Struct Mol Biol 32, 1006–1018. 10.1038/s41594-025-01491-y.

10. Shi, J., Hu, X., Guo, Y., Wang, L., Ji, J., Li, J., and Zhang, Z.R. (2019). A technique for delineating the unfolding requirements for substrate entry into retrotranslocons during endoplasmic reticulum-associated degradation. Journal of Biological Chemistry 294, 20084–20096. 10.1074/jbc.RA119.010019.

11. Tirosh, B., Furman, M.H., Tortorella, D., and Ploegh, H.L. (2003). Protein unfolding is not a prerequisite for endoplasmic reticulum-to-cytosol dislocation. Journal of Biological Chemistry 278, 6664–6672. 10.1074/jbc.M210158200.

12. Vitali, D.G., Fonseca, D., and Carvalho, P. (2024). The derlin Dfm1 couples retrotranslocation of a folded protein domain to its proteasomal degradation. Journal of Cell Biology 223. 10.1083/jcb.202308074.

13. Leto, D.E., Morgens, D.W., Zhang, L., Walczak, C.P., Elias, J.E., Bassik, M.C., and Kopito, R.R. (2019). Genome-wide CRISPR Analysis Identifies Substrate-Specific Conjugation Modules in ER-Associated Degradation. Mol Cell 73, 377–389.e11. 10.1016/j.molcel.2018.11.015.

14. Nejatfard, A., Wauer, N., Bhaduri, S., Conn, A., Gourkanti, S., Singh, N., Kuo, T., Kandel, R., Amaro, R.E., and Neal, S.E. (2021). Derlin rhomboid pseudoproteases employ substrate engagement and lipid distortion to enable the retrotranslocation of ERAD membrane substrates. Cell Rep 37. 10.1016/j.celrep.2021.109840.

15. Rao, B., Li, S., Yao, D., Wang, Q., Xia, Y., Jia, Y., Shen, Y., and Cao, Y. (2021). The cryo-EM structure of an ERAD protein channel formed by tetrameric human Derlin-1.

16. Schoebel, S., Mi, W., Stein, A., Ovchinnikov, S., Pavlovicz, R., DImaio, F., Baker, D., Chambers, M.G., Su, H., Li, D., et al. (2017). Cryo-EM structure of the protein-conducting ERAD channel Hrd1 in complex with Hrd3. Nature 548, 352–355. 10.1038/nature23314.

17. Guo, L., Liu, G., He, J., Jia, X., He, Y., Wang, Z., and Qian, H. (2025). Structural insights into the human HRD1 ubiquitin ligase complex. Nature Communications 16. 10.1038/s41467-025-61143-z.

18. Wu, X., Siggel, M., Ovchinnikov, S., Mi, W., Svetlov, V., Nudler, E., Liao, M., Hummer, G., and Rapoport, T.A. (2020). Structural basis of ER-associated protein degradation mediated by the Hrd1 ubiquitin ligase complex. Science 368. 10.1126/science.aaz2449.

19. Horn, S.C., Hanna, J., Hirsch, C., Volkwein, C., Schütz, A., Heinemann, U., Sommer, T., and Jarosch, E. (2009). Usa1 Functions as a Scaffold of the HRD-Ubiquitin Ligase. Mol Cell 36, 782–793. 10.1016/j.molcel.2009.10.015.

20. Hwang, J., Walczak, C.P., Shaler, T.A., Olzmann, J.A., Zhang, L., Elias, J.E., and Kopito, R.R. (2017). Characterization of protein complexes of the endoplasmicreticulum-associated degradation E3 ubiquitin ligase Hrd1. Journal of Biological Chemistry 292, 9104–9116. 10.1074/jbc.M117.785055.

21. Schulz, J., Avci, D., Queisser, M.A., Gutschmidt, A., Dreher, L.S., Fenech, E.J., Volkmar, N., Hayashi, Y., Hoppe, T., and Christianson, J.C. (2017). Conserved cytoplasmic domains promote Hrd1 ubiquitin ligase complex formation for ER-associated degradation (ERAD). J Cell Sci 130, 3322–3335. 10.1242/jcs.206847.

22. Belyy, V., Zuazo-Gaztelu, I., Alamban, A., Ashkenazi, A., and Walter, P. (2022). Endoplasmic reticulum stress activates human IRE1α through reversible assembly of inactive dimers into small oligomers. Elife 11. 10.7554/eLife.74342.

23. Sotolongo Bellón, J., Birkholz, O., Richter, C.P., Eull, F., Kenneweg, H., Wilmes, S., Rothbauer, U., You, C., Walter, M.R., Kurre, R., et al. (2022). Four-color single-molecule imaging with engineered tags resolves the molecular architecture of signaling complexes in the plasma membrane. Cell Reports Methods 2. 10.1016/j.crmeth.2022.100165.

24. Los, G. V., Encell, L.P., McDougall, M.G., Hartzell, D.D., Karassina, N., Zimprich, C., Wood, M.G., Learish, R., Ohana, R.F., Urh, M., et al. (2008). HaloTag: A novel protein labeling technology for cell imaging and protein analysis. ACS Chem Biol 3, 373–382. 10.1021/cb800025k.

25. Huang, C.H., Chu, Y.R., Ye, Y., and Chen, X. (2014). Role of HERP and a HERP-related protein in HRD1-dependent protein degradation at the endoplasmic reticulum. Journal of Biological Chemistry 289, 4444–4454. 10.1074/jbc.M113.519561.

26. Lin, L.L., Wang, H.H., Pederson, B., Wei, X., Torres, M., Lu, Y., Li, Z.J., Liu, X., Mao, H., Wang, H., et al. (2024). SEL1L-HRD1 interaction is required to form a functional HRD1 ERAD complex. Nat Commun 15. 10.1038/s41467-024-45633-0.

27. Grimm, J.B., Klein, T., Kopek, B.G., Shtengel, G., Hess, H.F., Sauer, M., and Lavis, L.D. (2016). Synthesis of a Far-Red Photoactivatable Silicon-Containing Rhodamine for Super-Resolution Microscopy. Angewandte Chemie 128, 1755–1759. 10.1002/ange.201509649.

28. Choi, H., Liao, Y.-C., Yoon, Y.J., Grimm, J., Lavis, L.D., Singer, R.H., and Lippincott-Schwartz, J. (2023). Lysosomal release of amino acids at ER three-way junctions regulates transmembrane 1 and secretory protein mRNA translation 2 3. bioRxiv. 10.1101/2023.08.01.551382.

29. Shibata, Y., Voeltz, G.K., and Rapoport, T.A. (2006). Rough Sheets and Smooth Tubules. Preprint at Elsevier B.V., https://doi.org/10.1016/j.cell.2006.07.019 10.1016/j.cell.2006.07.019.

30. Tinevez, J.Y., Perry, N., Schindelin, J., Hoopes, G.M., Reynolds, G.D., Laplantine, E., Bednarek, S.Y., Shorte, S.L., and Eliceiri, K.W. (2017). TrackMate: An open and extensible platform for single-particle tracking. Methods 115, 80–90. 10.1016/j.ymeth.2016.09.016.

31. Siggia, E.D., Lippincott-Schwartz, J., and Bekiranov, S. (2000). Diffusion in Inhomogeneous Media: Theory and Simulations Applied to Whole Cell Photobleach Recovery.

32. Nehls, S., Snapp, E.L., Cole, N.B., Zaal, K.J.M., Kenworthy, A.K., Roberts, T.H., Ellenberg, J., Presley, J.F., Siggia, E., and Lippincott-Schwartz, J. (2000). Dynamics and retention of misfolded proteins in native ER membranes.

33. Bi, C., Scrudders, K.L., Zheng, Y., Mahmoodi, M., Low-Nam, S.T., and Huang, F. (2025). SPTnet: a deep learning framework for end-to-end single-particle tracking and motion dynamics analysis. Preprint, https://doi.org/10.1101/2025.02.04.636521 10.1101/2025.02.04.636521.

34. Sun, Y., Yu, Z., Obara, C.J., Mittal, K., Lippincott-Schwartz, J., and Koslover, E.F. (2022). Unraveling trajectories of diffusive particles on networks. Phys Rev Res 4. 10.1103/PhysRevResearch.4.023182.

35. Snapp, E.L., Reinhart, G.A., Bogert, B.A., Lippincott-Schwartz, J., and Hegde, R.S. (2004). The organization of engaged and quiescent translocons in the endoplasmic reticulum of mammalian cells. Journal of Cell Biology 164, 997–1007. 10.1083/jcb.200312079.

36. Pisa, R., and Rapoport, T.A. (2022). Disulfide-crosslink analysis of the ubiquitin ligase Hrd1 complex during endoplasmic reticulum-associated protein degradation. Journal of Biological Chemistry 298. 10.1016/j.jbc.2022.102373.

37. Vasic, V., Denkert, N., Schmidt, C.C., Riedel, D., Stein, A., and Meinecke, M. (2020). Hrd1 forms the retrotranslocation pore regulated by auto-ubiquitination and binding of misfolded proteins. Preprint at Nature Research, https://doi.org/10.1038/s41556-020-0473-4 10.1038/s41556-020-0473-4.

38. Jumper, J., Evans, R., Pritzel, A., Green, T., Figurnov, M., Ronneberger, O., Tunyasuvunakool, K., Bates, R., Žídek, A., Potapenko, A., et al. (2021). Highly accurate protein structure prediction with AlphaFold. Nature 596, 583–589. 10.1038/s41586-021-03819-2.

39. Chambers, P., Pringle, C.R., and Easton, A.J. (1990). Heptad repeat sequences are located adjacent to hydrophobic regions in several types of virus fusion glycoproteins.

40. Peterson, B.G., Hwang, J., Russ, J.E., Schroeder, J.W., Freddolino, P.L., and Baldridge, R.D. (2023). Deep mutational scanning highlights a role for cytosolic regions in Hrd1 function. Cell Rep 42. 10.1016/j.celrep.2023.113451.

41. Moochickal Assainar, B., Ragunathan, K., and Baldridge, R.D. (2024). Direct observation of autoubiquitination for an integral membrane ubiquitin ligase in ERAD. Nat Commun 15. 10.1038/s41467-024-45541-3.

