## Supplementary Figures for "Stable Hrd1 tetramers at the heart of the retrotranslocon in living cells"

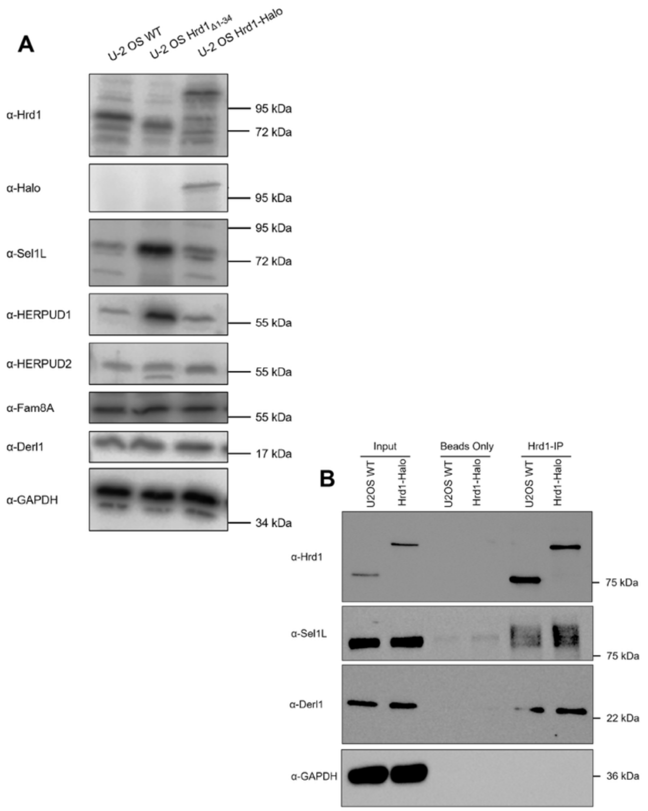


**Supplementary Figure 1: Hrd1 complex composition is unaltered upon HaloTag integration.**

(A) Western blot of cell lysates from genetically modified U-2 OS cell lines detecting all major Hrd1 complex components. Note that the abundance of non-Hrd1 complex components is similar between the WT and Hrd1-HaloTag lines, in contrast to the line expressing a nonfunctional version of Hrd1, which shows elevated levels of Sel1L and HERPUD1 as expected in response to requisite ER stress.

(B) Immunoprecipitation of WT and Hrd1-Halo cells precipitates Sel1L and Derlin1. There were no detectable differences in pull-down behavior between WT and Hrd1-Halo.

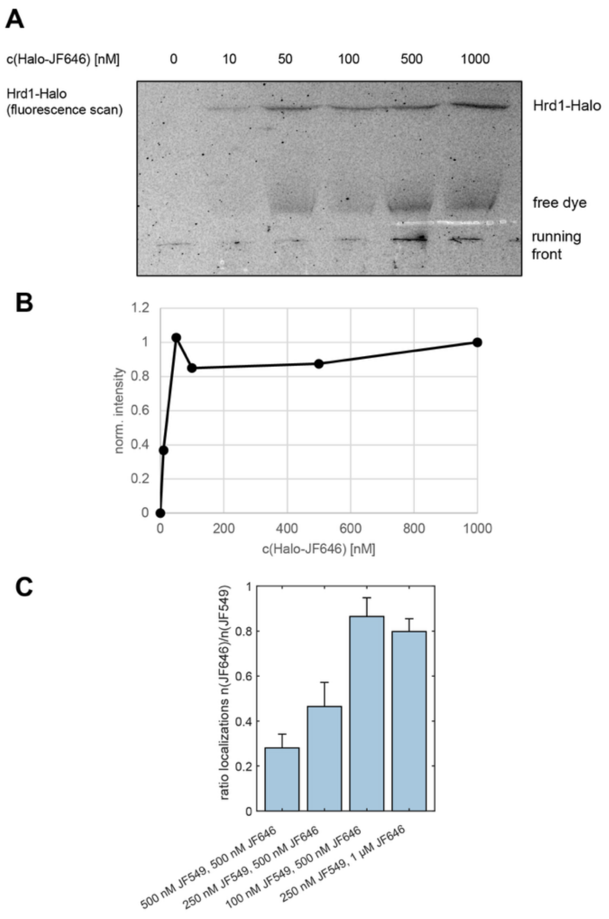


**Supplementary Figure 2: Optimization of Hrd1-Halo labeling, showing saturation of HaloTag ligand at ~50 nM.**

(A) Fluorescence scan SDS-PAGE separating U-2 OS Hrd1-Halo cell lysates previously labeled with the indicated concentrations of JF646-Halo for 10 minutes.

(B) Quantification of A normalized on the maximal value. Labeling reaches saturation at 50 nM.

(C) Number of localized emitters in dual color TIRF microscopy of U-2 OS Hrd1-Halo cells labeled for 10 minutes with the indicated concentrations. n(cells) = 5.

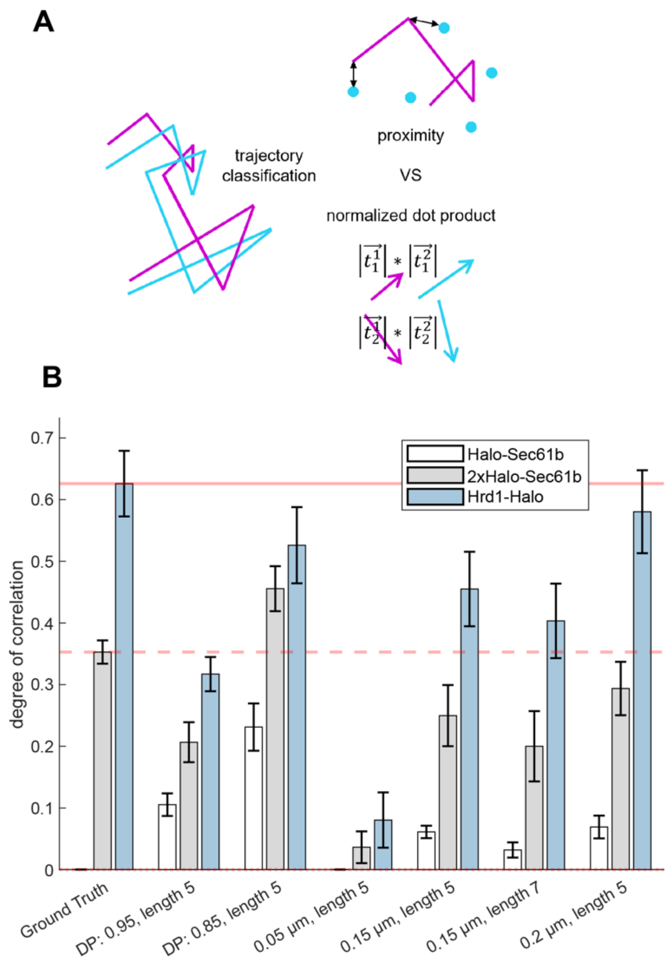


**Supplementary Figure 3: Comparison of direction and proximity-based trajectory classification.**

(A) Schematic representation of the two classification approaches tested. Proximity based correlation measures the closest distance of any point in a trajectory to an emitter in the second channel. Prolonged proximity below a cut-off value leads to a trajectory to be classified as correlated. Directionality based correlation compares the direction and length of steps from trajectories from both channels by computing their normalized dot-product. If trajectories show prolonged normalized dot-products above a cut-off value the trajectories are registered as correlated.

(B) Degree of correlation of Halo-Sec61b, 2xHalo-Sec61b and Hrd1-Halo as estimated by the correlation classifier in comparison to a manually classified ground truth (n=5 cells). The classifiers were tested using different cut-off values for the normalized dot-product (DP), required length of correlation and distance. 150 nm and 7 correlated steps were chosen as a compromise between high identification rate and low false positive rate.

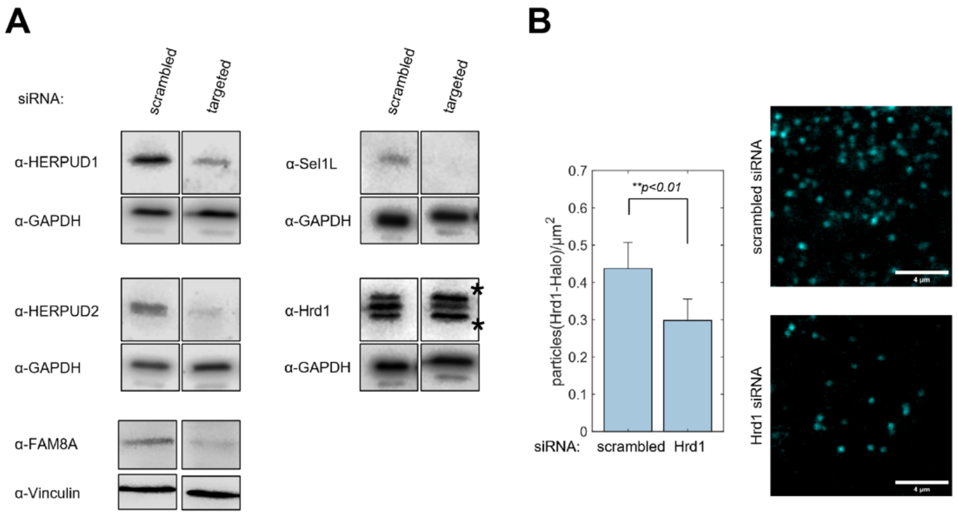


**Supplementary Figure 4: Validation of siRNA mediated knockdown of Hrd1 complex components.**

(A) Representative western blots of U-2 OS cell lysates after 48 h of treatment with the indicated siRNA pools.

(B) Validation of partial Hrd1 knockdown by Hrd1-Halo single-molecule counting using TIRF microscopy. siRNA leads to a significant reduction of detected emitters. Significance was tested using a one-sided ANOVA. n(cells) = 127,52.

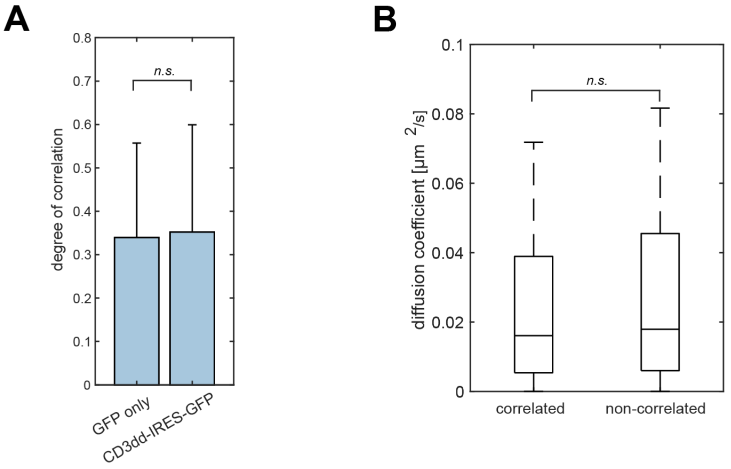


**Supplementary Figure 5: Hrd1 complex formation is not sensitive to over expression of obligate ERAD cargo.**

(A) Degree of autocorrelation in Hrd1 dcSMT in dependence of substrate overexpression. CD3dd-HA had no impact on Hrd1 correlation. Significance was tested using a one-sided ANOVA. n(cells) = 27,26 n(trajectories) = 559,391.

(B) Comparison of diffusion coefficients of correlated and non-correlated Hrd1-Halo particles. Significance was tested using a one-sided ANOVA. n(correlated/non-correlated) = 1527.

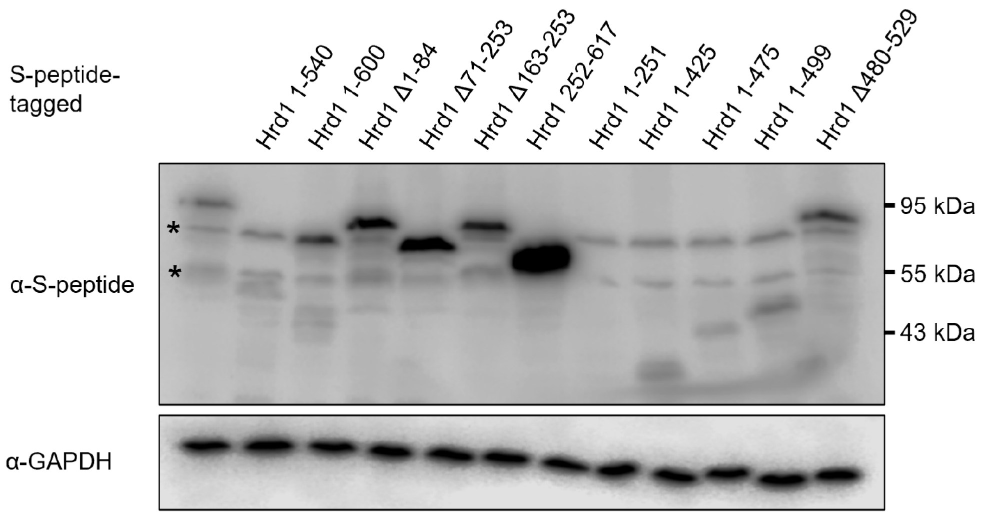


**Supplementary Figure 6: Confirmation of overexpression of S-peptide tagged Hrd1.** Western blot of cell lysates expressing the S-peptide tagged Hrd1 variants used in the competitive dcSMT assay after 24 h of expression. Similar as observed in Schulz et al., Hrd1 1-251 could not be detected by Western blot.

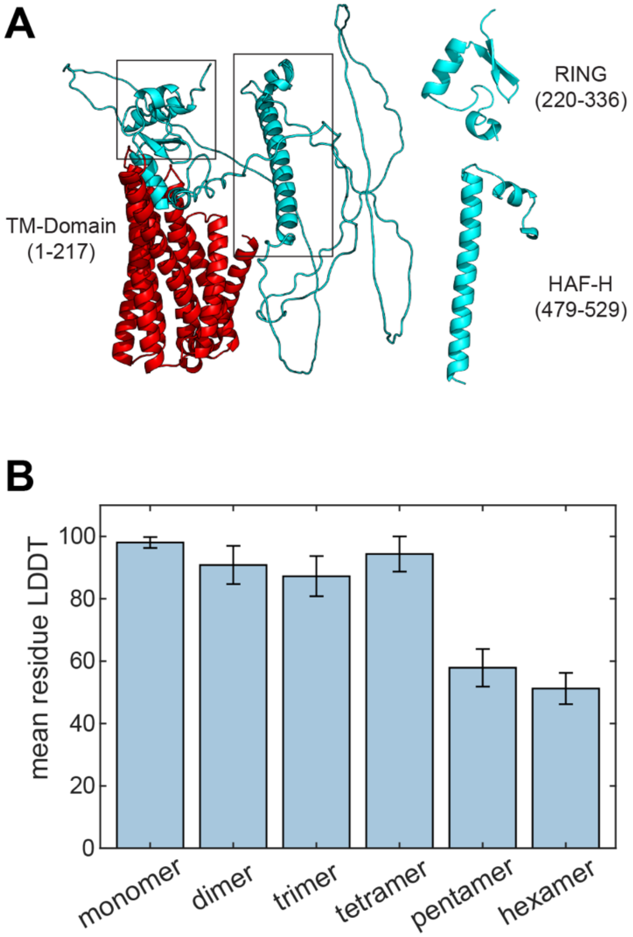


**Supplementary Figure 7: AlphaFold prediction of Hrd1.**

(A) Prediction of the structure of monomeric full-length Hrd1.

(B) Comparison of average, residue-wise pLDDT scores for different copy numbers of Hrd1480-529. Up to four copies lead to acceptable scores.

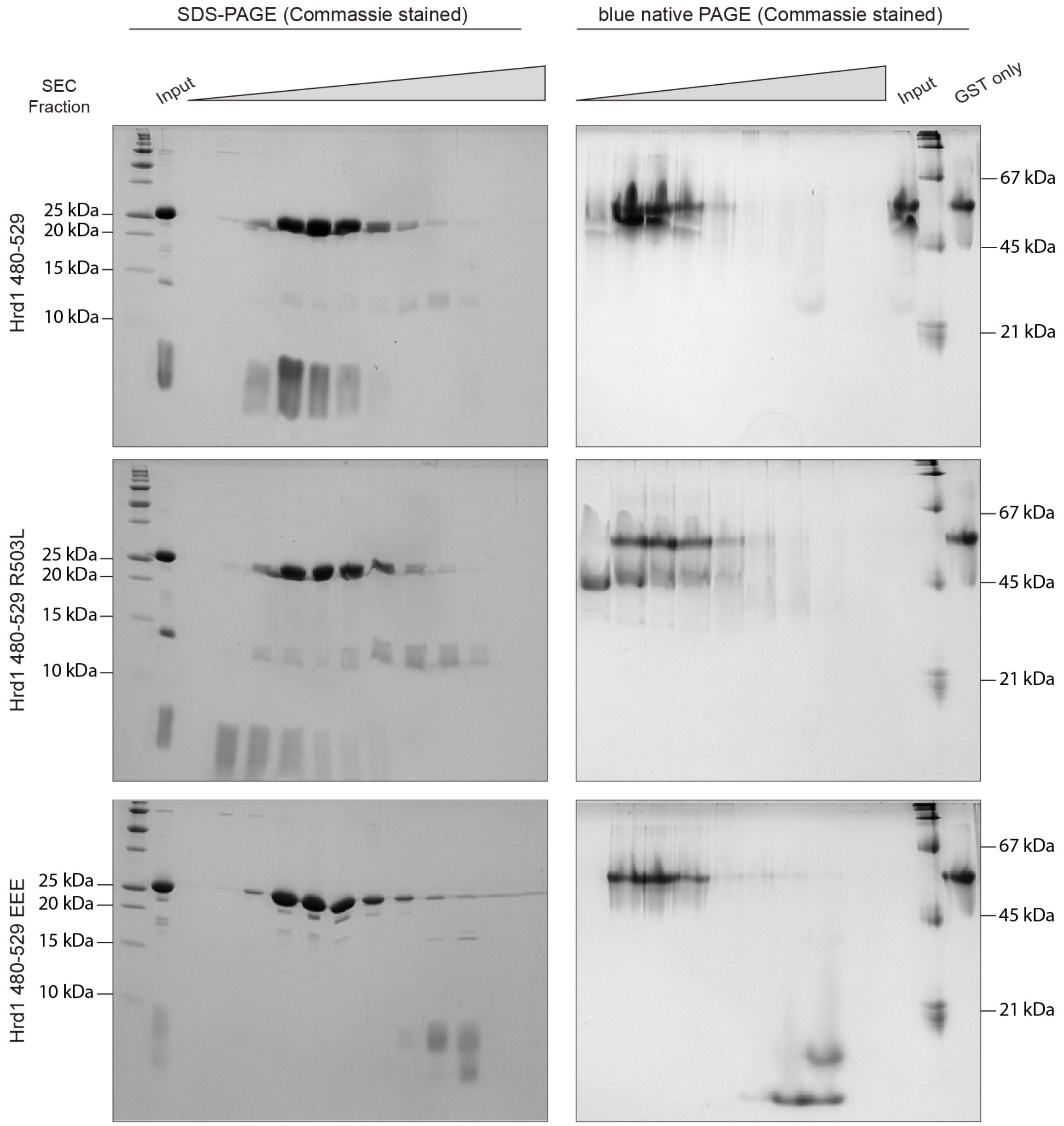


**Supplementary Figure 8: Hrd1 480-529 migrates as a species with approximately 4 times its apparent molecular weight under native conditions.** Replacing amino acids participating in the heptad-repeats leads to behavior in line with a monomer. Commassie stained SDS- and blue native PAGE of SEC fractions of purified Hrd1 480-529 variants. All samples contain cleaved GST-tag.

<table>
<tr><th>Fraction</th><th>hSyvn_wt<br>⊠ 5420.19 Da</th><th>hSyvn_RL<br>⊠ 5377.16 Da</th><th>hSyvn_EEE<br>⊠ 5468.06 Da</th></tr>
<tr>
<td>loaded onto SEC</td>
<td>on SDS gel:<br>mixture of GST, wt protein & E.coli contaminant
<table>
<tr><td>4031.85<br>(16.5%)</td><td>degradation product of hSyvn_wt[1]</td></tr>
<tr><td>5420.61<br>(100%)</td><td>hSyvn_wt<br>(10840.42 Da with 4.25% deconvolution artefact thereof)</td></tr>
<tr><td>26431<br>(2.5 %)</td><td>GST-Tag</td></tr>
<tr><td>14829.30<br>(0.1 %)</td><td>E.coli protein*<br>(as very tiny peak next to other small peaks detectable)</td></tr>
</table>
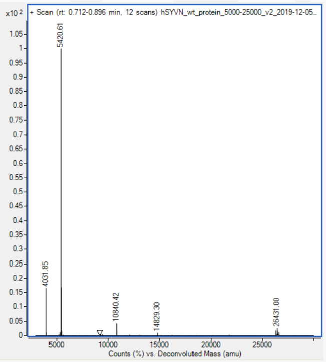

[1]Degradation<br>5421-4032=-1389 Da<br>GPLGSEELRALEGHERQHLEALLQSLRNIHTLLDAA~~MLQINQYLTVLA~~</td>
<td>on SDS gel:<br>mixture of GST, RL protein & E.coli contaminant
<table>
<tr><td>5377.41<br>(100%)</td><td>hSyvn_RL<br>(10754.37 Da with 5.9% deconvolution artefact thereof)</td></tr>
<tr><td>26431.10<br>(2.7 %)</td><td>GST-Tag</td></tr>
<tr><td>14829.44<br>(3.5%)</td><td>E.coli protein*<br>(as very tiny peak next to other small peaks detectable)</td></tr>
</table>
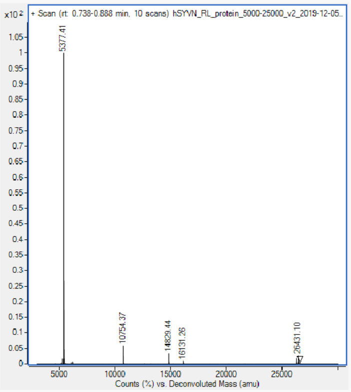
</td>
<td>on SDS gel:<br>mixture of GST, EEE protein, some other degradation bands (between 18-24 kDa)
<table>
<tr><td>4047.71<br>(100 %)</td><td>degradation product of hSyvn_EEE[2]</td></tr>
<tr><td>5468.66<br>(34%)</td><td>hSyvn_EEE</td></tr>
<tr><td>26430.81<br>(6 %)</td><td>GST-Tag</td></tr>
</table>
⊠ no 14829 Da E.coli contamination detectable (and also not visible on gel)
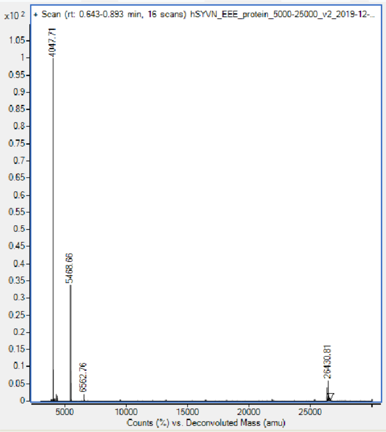

[2]Degradation<br>5469-4048=-1421 Da<br>GPLGSEELRALEGHERQHLEARLQSLRNIHTLEDAA~~MLQENQYLTVEA~~</td>
</tr>
</table>

**Supplementary Figure 9: The purified samples contain only the Hrd1 fragment and cleaved GST-tag.** Intact-mass MS of the SEC input sample shown in (S8). According to the detected molecular weights all samples exclusively contain the Hrd1 fragment, cleaved GST-tag and a minimal contamination with bacterial derived protein.
